# Effects of two centuries of global environmental variation on phenology and physiology of *Arabidopsis thaliana*

**DOI:** 10.1101/424242

**Authors:** Victoria L. DeLeo, Duncan N. L. Menge, Ephraim M. Hanks, Thomas E. Juenger, Jesse R. Lasky

## Abstract

Intraspecific trait variation is caused by genetic and plastic responses to environment. This intraspecific diversity is captured in immense natural history collections, giving us a window into trait variation across continents and through centuries of environmental shifts. Here we tested if hypotheses based on life history and the leaf economics spectrum explain intraspecific trait changes across global spatiotemporal environmental gradients. We measured phenotypes on a 216-year time series of *Arabidopsis thaliana* accessions from across the native range and applied spatially varying coefficient models to quantify region-specific trends in trait coordination and trait responses to climate gradients. All traits exhibited significant change across space and/or through time. For example, δ^15^N decreased over time across much of the range and leaf C:N increased, consistent with predictions based on anthropogenic changes in land use and atmosphere. Plants were collected later in the growing season in more recent years in many regions, possibly because populations shifted toward more spring germination and summer flowering as opposed to fall germination and spring flowering. When climate variables were considered, collection dates were earlier in warmer years, while summer rainfall had opposing associations with collection date depending on regions. There was only a modest correlation among traits, indicating a lack of a single life history/physiology axis. Nevertheless, leaf C:N was low for summer- versus spring-collected plants, consistent with a life history-physiology axis from slow-growing winter annuals to fast-growing spring/summer annuals. Regional heterogeneity in phenotype trends indicates complex responses to spatiotemporal environmental gradients potentially due to geographic genetic variation and climate interactions with other aspects of environment. Our study demonstrates how natural history collections can be used to broadly characterize trait responses to environment, revealing heterogeneity in response to anthropogenic change.

## Introduction

An organism’s fitness is determined by the interaction between its traits and its environment. Organisms respond to environmental gradients in diverse ways, including genetic and plastic shifts in life history, phenology, and physiology (Burghardt, Metcalf, Wilczek, Schmitt, & Donohue, 2015; Reich, 2014; Wright et al., 2004). Spatial and temporal environmental gradients can promote phenotypic plasticity or generate varying selection across which populations adapt to local conditions (Bradshaw, 1965; Henn et al., 2018; Joshi et al., 2001; Leimu & Fischer, 2008; Linhart & Grant, 2002; Matesanz, Gianoli, & Valladares, 2010; Turesson, 1922). By studying how phenology and physiology change across environments through space and time we can learn about mechanisms of adaptive environmental response and biological constraints.

Anthropogenic global change has led to dramatic phenotypic changes in many organisms. For example, many species are shifting their ranges poleward and temperate spring phenology is advancing (Parmesan & Yohe, 2003). However, anthropogenic global change is multi-faceted, involving climate, nitrogen deposition, atmospheric CO_2_, and land use. Our understanding of the specific environmental drivers of phenotypic change has been hampered both by insufficient long-term datasets and by the complexities of interacting and correlated environmental variables. Furthermore, many populations and species do not exhibit the stereotypic advancing temperate phenology and poleward range shifts (Both et al., 2004; CaraDonna, Iler, & Inouye, 2014; Park et al., 2018). These diverse responses can be caused by geographic variation in the rate of environmental change or by intraspecific genetic variation, clouding our understanding of anthropogenic impacts. To address these challenges, we collected physiology and phenology data for *Arabidopsis thaliana* (hereafter, *Arabidopsis*) specimens over the last 200 years and across its native range, and we tested relationships between climate and traits using spatial generalized additive models (GAMs) to account for geographic structure in environmental response.

*Arabidopsis* is a powerful system for studying phenotypic change across space and climate gradients. Past studies have found that *Arabidopsis* populations exhibit genetic differences among populations likely due to isolation by distance (Alonso-Blanco et al., 2016; Horton et al., 2012; Ostrowski et al., 2006; Platt et al., 2010) and local adaptation (Fournier-Level et al. 2011, Hancock et al. 2011, Lasky et al. 2012). Genetic differences in flowering time among populations may be due to local adaptation (Atwell et al., 2010; Burghardt et al., 2015; Tabas-Madrid et al., 2018), with northern genotypes having later flowering times (Atwell et al., 2010; Stinchcombe et al., 2004). *Arabidopsis* physiology also shows evidence of a role in local adaptation. Genotypes from regions of greater precipitation have faster growth and lower leaf vein density, and genotypes from colder temperatures have increased leaf thickness and wider leaf minor vein cross section (Adams, Stewart, Cohu, Muller, & Demmig-Adams, 2016; Sack et al., 2012; Sartori et al., 2018). These findings provide a lens through which to interpret phenotypic variation among plants in nature. In turn, museum collections offer broadly distributed sampling in space and time to test the relationships between phenotypes and environment in nature (Lang, Willems, Scheepens, Burbano, & Bossdorf, 2018; Willis et al., 2017).

Multiple frameworks of plant life history and physiology variation correspond to a continuum of fast to slow life histories (e.g. Grime, 1977; Westoby, 1998; I. J. Wright et al., 2004). Individuals that use a fast strategy are characterized by fast relative growth, early reproduction, and intensive use of nutrients or water, while individuals with a slow strategy are characterized by slow growth, late reproduction, and more measured use of nutrients and water. The Leaf Economics Spectrum (LES) hypothesizes that leaves vary along single life history-physiology axis from fast to slow in association with large-scale climate gradients (Reich, 2014; Reich et al., 2003; Wright et al., 2004). The LES predicts that lower nitrogen concentration leaves (high C:N, low proportion N) are found in drier and in hotter areas, possibly in part because of investment in non-photosynthetic leaf features, e.g. veins (Blonder, Violle, Bentley, & Enquist, 2011; Easlon et al., 2014; Sack et al., 2012) (see Table 1 for phenotype/environment hypotheses and how they relate to the fast/slow framework). Low N leaves are thicker (high mass to area) and provide protection against stress (drought) at the expense of a N investment in photosynthesis, resulting in a slower life cycle (Evans, 1989; Stocking & Ongun, 1962). Although community-wide turnover in mean traits across natural environments often follows LES predictions, *within* species trait variation often defies LES predictions (Anderegg et al., 2018; Hu et al., 2015; J. P. Wright & Sutton-Grier, 2012). Nevertheless, *Arabidopsis* exhibits genetic variation in traits that corresponds to LES predictions (Easlon et al., 2014; Sartori et al., 2018; Vasseur, Violle, Enquist, Granier, & Vile, 2012); individuals with rapid life histories have physiological traits tied to fast growth and resource acquisition (e.g. high stomatal conductance, high leaf area relative mass (Specific Leaf Area, or SLA)) (Lovell et al., 2013; McKay, Richards, & Mitchell-Olds, 2003; Sartori et al., 2018; Wolfe & Tonsor, 2014).

**Table 1:** Hypothesized responses of phenotypes to increases in temperature, rainfall, or year, or how traits would change along a faster life history strategy. Year trends are predicted due to elevated CO_2_, nitrogen deposition, or elevated temperatures. Citations for hypotheses: (Amundson et al., 2003^1^; BassiriRad et al., 2003^2^; Burghardt et al., 2015^3^; Craine et al., 2009^4^; Diefendorf et al., 2010^5^; Drake, Hanson, Lowrey, & Sharp, 2017^6^; Gill et al., 2002^7^; McLauchlan, Ferguson, Wilson, Ocheltree, & Craine, 2010^8^; Menzel et al., 2006^9^; Ordoñez et al., 2009^10^; Peñuelas et al., 2004^11^; Reich, Hungate, & Luo, 2006^12^; Seibt, Rajabi, Griffiths, & Berry, 2008^13^; Sparks & Carey, 2006^14^; Stock & Evans, 2006^15^; I. J. Wright et al., 2004^16^)

|  | Temperature | Rainfall | Year | Fast Life History |
| --- | --- | --- | --- | --- |
| $\Delta^{13}\text{C}$ | +<sup>13</sup> | +<sup>5</sup> | +<sup>6</sup> | + |
| $\delta^{15}\text{N}$ | +<sup>1,4</sup> | -<sup>1,4</sup> | -<sup>2,8,15</sup> | No change |
| C:N | -<sup>10</sup> in cold regions<br>+<sup>16</sup> in warm regions | -<sup>16</sup> | +<sup>7,8,12</sup> | - |
| Photothermal Units | + or no change | + | No change | - |
| Collection Date | -<sup>3,9</sup> | +<sup>11</sup> | -<sup>9,14</sup> | - |

Isotopic signatures provide clues to organismal life histories. In plants, Δ^13^C measures discrimination against ^13^C in photosynthesis and is an indicator of pCO_2_ within leaves (C_i_) relative to atmospheric pCO_2_ (C_a_) (Farquhar, O’Leary, & Berry, 1982). C_i_ declines when stomata are closed, which may be a conservative life history response to soil drying, while C_a_ declines with elevation. Thus, we expect Δ^13^C to increase in moist growing environments and decrease with elevation (Diefendorf, Mueller, Wing, Koch, & Freeman, 2010; Farquhar et al., 1982; Zhu, Siegwolf, Durka, & Körner, 2010) (Table 1). δ^15^N (the ratio of ^15^N to ^14^N) can be affected by nitrogen allocation and so might reflect variation in C:N (Stock & Evans, 2006). However, variation in leaf N and δ^15^N may also directly reflect changing environments (N deposition, biogeochemical cycling, e.g. Pardo *et al*. 2007) rather than plant traits, with both increasing with temperature and leaf N increasing and δ^15^N decreasing with rainfall (Table 1).

Fast/slow strategies also correspond to phenological variation, especially in annual plants. In seasonal environments, phenology is constrained by seasonality and simultaneously determines the environment encountered during vulnerable stages. *Arabidopsis* development can be highly sensitive to moisture, temperature, and photoperiod (Burghardt et al., 2015; Wilczek et al., 2009). For example, although warmth can increase growth rates, many *Arabidopsis* genotypes require winter cold cues (known as vernalization) to transition to spring flowering. Fast life histories can allow spring or summer annual life cycles, where a plant germinates and flowers within a single season, while slow life histories and vernalization requirements result in a winter annual cycle, where a plant germinates in the fall and flowers the following spring. Rapid development and reproduction can allow *Arabidopsis* plants to escape drought (McKay et al., 2003), while slower flowering plants can exhibit drought avoidance strategies of minimizing water loss (e.g. through stomatal closure) or maximizing water uptake (Kenney, Mckay, Richards, & Juenger, 2014; Ludlow, 1989). Because herbarium specimens are typically reproductive, it is challenging to infer germination times based on collection dates. However, information on physiology and climate preceding collection may provide information on life history variation.

Standardized metrics facilitate the comparison of ecologically relevant phenological variation among sites that differ in climate and seasonal timing. Photothermal units (PTUs) integrate developmental time under favorable temperatures and light and account for much of the environmental influence on flowering dates in *Arabidopsis* (Brachi et al., 2010; Wilczek et al., 2009). In essence, PTUs estimate how far along in a growing season an event occurs. Measures of developmental time standardized to environmental conditions can better capture genetic variation in development compared to raw flowering dates in *Arabidopsis*, the latter of which are strongly driven by environment (Brachi et al., 2010). Without knowing an exact germination date, it is impossible to perfectly approximate the climate experienced in the wild, yet even a rough estimate using an arbitrary date allows us to compare changes in the climate experienced at flowering. Variation in herbarium collection dates is a reliable proxy for variation in phenology of flowering date (Davis, Willis, Connolly, Kelly, & Ellison, 2015; MacGillivray, Hudson, & Lowe, 2010; Miller-Rushing, Primack, Primack, & Mukunda, 2006). In addition, low PTUs at collection hints at a winter annual growing pattern (because growth is occurring before PTU calculations begin), so we can use this measurement to search for regional variation in life history. Combined with existing knowledge the phenology of ecotypes from different sites, PTUs may help reveal phenological adaptation along spatial and temporal environmental gradients.

Here, we leverage the immense fieldwork underlying natural history collections to investigate how intraspecific diversity is structured through time and along spatiotemporal climate gradients. Specifically, we use thousands of *Arabidopsis* specimens that span over 200 years of sampling across *Arabidopsis*’ range in Eurasia and Northern Africa. We quantify the spatial patterns of *Arabidopsis*’ phenotypic variation along environmental gradients, which allows us to put temporal trends in context. We hypothesized that for natural *Arabidopsis* populations, phenotype-environment correlations would follow fast-slow predictions of LES and phenology traits (Table 1). We combine these records with global gridded climate data to ask three questions about *Arabidopsis* in nature:

1. To what degree does intraspecific trait variation among wild individuals fall along a single coordinated life history-physiology axis?
2. Do life history and physiology vary across spatial environmental gradients in long-term average conditions, suggesting adaptive responses consistent with the LES?
3. Have life history and physiology changed over the last two centuries? In particular, have changes tracked climate fluctuations, suggesting adaptive responses consistent with the LES?

## Materials & Methods

### Samples

Our set of samples (N= 3443) included *Arabidopsis thaliana* herbarium and germplasm accessions with known collection dates between 1794 and 2010 from the native range of *Arabidopsis* in Europe, the Middle East, Central Asia, and North Africa (Hoffmann, 2002). Wild-collected germplasm accessions with known collection date and location (N = 447) were included only in models of phenology. Information on germplasm accessions came from the *Arabidopsis* Biological Resource Center (https://abrc.osu.edu/). For each herbarium specimen (N=2663) we visually verified species identification and reproductive status as simultaneously flowering and fruiting. Samples that were only fruited/senesced, only flowering, or had neither open flowers nor fruits were excluded to focus on a relatively uniform developmental stage (see Supplementary Table 1). This consistency is important for assessing C:N, since progression of plant development involves reallocation of nutrients, and for a meaningful characterization of phenology with collection date (Himelblau & Amasino, 2001). Furthermore, too few samples (178/2663) were in other phenological stages to allow for a rigorous comparison. We excluded dozens of misidentified specimens, highlighting the importance of verification of information in natural history collections (cf. unverified data in some online databases). Samples with too low precision in collection date (less precise than a single month) were excluded from phenological analysis.

### Leaf traits

To test LES hypotheses for response to environment, we measured Δ ^13^C, δ^15^N, and C:N from leaf tissue of herbarium samples. We removed and pulverized leaf samples (mean weight = 2.75 mg) of a subset of our quality-checked herbarium specimens and sent them to the UC-Davis Stable Isotope Facility. In total, we obtained values for δ^15^N, δ^13^C, C:N, and proportion N in 459 accessions, although 5 samples failed for δ^13^C and 1 sample was missing failed for both C:N and proportion N values.

We measured leaf δ^13^C (isotope ratio), but atmospheric δ^13^C has changed dramatically over the time period of this study due to fossil fuel emissions. Thus we converted leaf isotope ratio (δ^13^C) to discrimination (Δ^13^C) using an estimate of the atmospheric δ^13^C time series (McCarroll & Loader, 2004) from 1850 to 2000, continuing linear extrapolation beyond 2000, using the 1850 value for earlier specimens, and the equation of Farquhar et al. (1989), 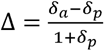, where δ*_a_* is the isotope ratio in the atmosphere and δ*_p_*is the isotope ratio in plant tissue (ratios relative to a standard).

### Phenology

To estimate accumulated photothermal units (PTU) at date of collection, we used the equation of Burghardt et al. (2015) to model the hourly temperature values for the accumulation of sunlight degree hours between January 1 and dusk on the date of collection at each accession’s coordinate. Daylength was approximated with the R package geosphere (Hijmans, 2017). Monthly temperature values for the period 1900-2010 came from the Climate Research Unit time series dataset v4.01 (Harris, Jones, Osborn, & Lister, 2014). PTUs were only calculated for specimens collected after 1901 (N = 2488) due to the historical limit of the monthly temperature data. Daily temperatures were interpolated from monthly temperatures using the function splinefun in R on the “periodic” setting. PTUs calculated from January 1^st^ will not completely account for the climate experienced by plants that germinate in the fall. However, for the same developmental time (PTUs), determined by weather conditions, winter annuals are expected to flower earlier in a growing season compared to spring annuals. Comparing changes in PTU at collection to changes in date of collection might provide clues as to where climate is driving flowering time shifts and where flowering time is responding to pressures other than temperature.

### Statistical analysis

*Arabidopsis* displays substantial genetic diversity in environmental response between genotypes from different regions (e.g. Exposito-Alonso et al., 2018; Lasky, Forester, & Reimherr, 2018). Thus, we employed a regression model with spatially varying coefficients (generalized additive models, GAMs) to account for regional differences in responses to environment, much of which may have a genetic component (Wheeler & Waller, 2009; Wood, 2006). GAMs allow fitting of parameters that vary smoothly in space (i.e. parameter surfaces) and can thus capture spatially varying relationships between predictors and the response of interest, such as we see in ecological processes (Yee & Mackenzie, 2002; Yee & Mitchell, 2006). The spatially varying coefficients fit by GAM allow us to infer from the data where relationships between variables change, as opposed to binning data into a set of fixed (and possibly artificially defined) regions. In a standard linear statistical model, the effect of a covariate x (i.e., the effect of x = January Minimum Temperature) at site i is a linear function of the covariate: x_i_*β. Note that differences across sites in x_i_*β are completely controlled by differences in the covariate x at different sites.

In the spatially-varying GAMs we consider in this study, we allow the effect of covariates to vary across space, with the effect of a covariate x at location i being x_i_*β_i_ where β_i_ is the linear effect of x at the i^th^ spatial location. That is, we model allow the effect of the covariate x to vary across space. Thus, the effect of (for example) January Minimum Temperature might be different in Europe than it would be in Southeast Asia.

It would be impossible to uniquely identify a completely different effect β_i_ at every location, as we only have one replicate at each site, and there would be no replication. Spatially-varying GAMs address this identifiability by smoothing β_i_ across space and requiring that effects β_i_ and β_j_ for sites i and j that are close in space be very similar to each other. The degree of spatial smoothness in the GAM is chosen by cross-validation to ensure that the GAM does not overfit the data. Thus, spatially-varying coefficient models provide an approach for modeling variability in the effect of covariates across space, with the effect constrained to vary slowly across space to ensure identifiability and combat overfitting.

Each cell in our 140×200 grid model rasters corresponded to 53.1 km East/West at the lowest latitude (28.16°, versus 20.3 km at 68.18° N) and 31.8 km North/South (calculated using Vincenty ellipsoid distances in the geosphere package). Smaller grid cells allow for more finely smoothed slope values. Model predictions farther than 200km from a sampled accession were discarded when visualizing results.

We selected climate variables based on knowledge of critical *Arabidopsis* developmental times and likely environmental stressors: average temperature in April, when warmth is expected to accelerate development (AprilMean in the models), minimum temperature in January, when vernalization cues are likely accumulating or when Mediterranean plants are in early growth (JanMinimum), and July aridity index (AI), when summer drought may be most likely (Fournier-Level et al., 2013; Hoffmann, 2002; Lasky et al., 2012; Wilczek, Cooper, Korves, & Schmitt, 2014). Our analyses should not be highly sensitive to the exact calendar month chosen, given the high correlation in conditions between consecutive months (e.g. warm Aprils tend to be followed by warm Mays). Aridity index was calculated from July precipitation divided by July potential evapotranspiration (PET) (United Nations Environment Program, 1997). These climate gradients were generally not strongly correlated (July Aridity to April Mean Temperature r = −0.33; July Aridity to January Minimum Temperature r = −0.12; January Minimum Temperature to April Mean Temperature r = 0.71 by Pearson’s product-moment correlation). We took temperature, precipitation, and PET values from the Climate Research Unit time series dataset, using values for the year of collection (New, Hulme, & Jones, 2000).

First, to study trait correlations that might indicate a fast-slow life history (Question 1), we performed a Principal Components Analysis of flowering time, Δ^13^C, δ^15^N, and C:N ratio and tested pairwise associations between traits. We considered how traits co-vary by calculating the Pearson’s correlation coefficients between traits and by Principal Components Analysis. We also fit GAMs (described in detail below) with spatially varying intercepts allowing measured phenotypes as both response and predictor variables to observe how the correspondence of traits changes through space.

Second, to study phenotypic responses to spatial gradients in long-term average climates (Question 2), we fit models with spatially varying coefficients for long-term, 50-year climate averages at each location (“spatial climate models”, Equation 1). We scaled these climate covariates and year of collection to unit standard deviation (Hijmans, Cameron, Parra, Jones, & Jarvis, 2005). In these models of responses to long-term average conditions, year of collection can be considered a nuisance variable, accounting for temporal variation at a location that may be important but is not the focus of this specific model. In spatial climate models, we used a single global intercept. Spatial climate models included specimens from all years with phenotype data.

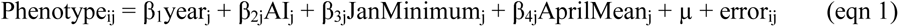

In all models, the subscript j denotes location and i denotes year of collection. For the temporal climate anomaly model, the spatially varying intercept is denoted by µ_j_, where the “j” subscript indicates that the intercept varies with location. The errors are assumed to be independent, be normally distributed, and have constant variance.

Next, to assess how phenotypes have changed across the last two centuries (Question 3), we tested a model with spatially-varying coefficients for the effect of year, allowing for geographic variation in temporal trends (hereafter, “year models”). The model also included spatially varying intercepts to account for regional differences in long-term mean phenotypes. Year models included all specimens with data for a particular phenotype.

Finally, to assess how temporal fluctuations in climate drive phenotypic change (Question 3), we fit models with the three climate covariates for the year of collection (Equation 2). We converted climate covariates to local anomalies by standardizing them relative to the entire time-series for a given grid cell to unit standard deviation and mean zero (“temporal climate anomaly models”). In standardizing climate fluctuations to the climate record of a location, we assume that the effect of an anomaly on the response variable is best captured by the relative strength (and direction) of an anomaly (relative to an average anomaly) rather than how extreme an anomaly is in general.

The model also included spatially varying intercepts to account for regional differences in long-term mean phenotypes. Temporal climate models only included specimens from after 1900, when we had data on monthly climate from CRU. These models had the following structure

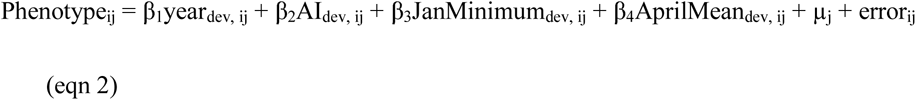

Models were fit in R (version 3.5.0, R Core Team 2011) using the ‘gam’ function in package mgcv (version 1.8-17, Wood 2011). We allowed the model fitting to penalize covariates to 0 so that covariates weakly associated with phenotypes could be completely removed from the model; thus, using the mgcv package we can achieve model selection through joint penalization of multiple model terms. Coefficients in spatially varying coefficient models represent the relationship between each term and phenotype at each geographic point (indexed by *j* in our models).

We considered two other spatially-varying environmental variables of interest: elevation and N deposition. We left elevation and nitrogen deposition covariates out of the final models because inclusion resulted in instability in the numerical routines the GAM software (mgcv) used to estimate parameters and approximate Hessian matrices needed for confidence intervals. See supplemental material for more information on these covariates. Including only the variables of the three climate covariates and year resulted in numerically stable estimates. In addition, scaling of year and climate variables tended to reduce the concurvity of variables and increase stability.

Code for all the models and plots will be included as a supplement and will be available on github.

## Results

### Distribution of samples through time and space

Samples were broadly distributed, with dense collections in Norway/Sweden, the Netherlands, and Spain (reflecting major herbaria used in the study), and sparser collections to the east (Figure 1). The subset of samples with tissue analysis spanned the extent of the geographic distribution of all samples. Tissue sampling was most dense in Norway, Spain, and the UK and sparser elsewhere in the range. The earliest collection date we used was 1794, but a greater number of samples were available from the 1900s onwards.

**Figure 1:**
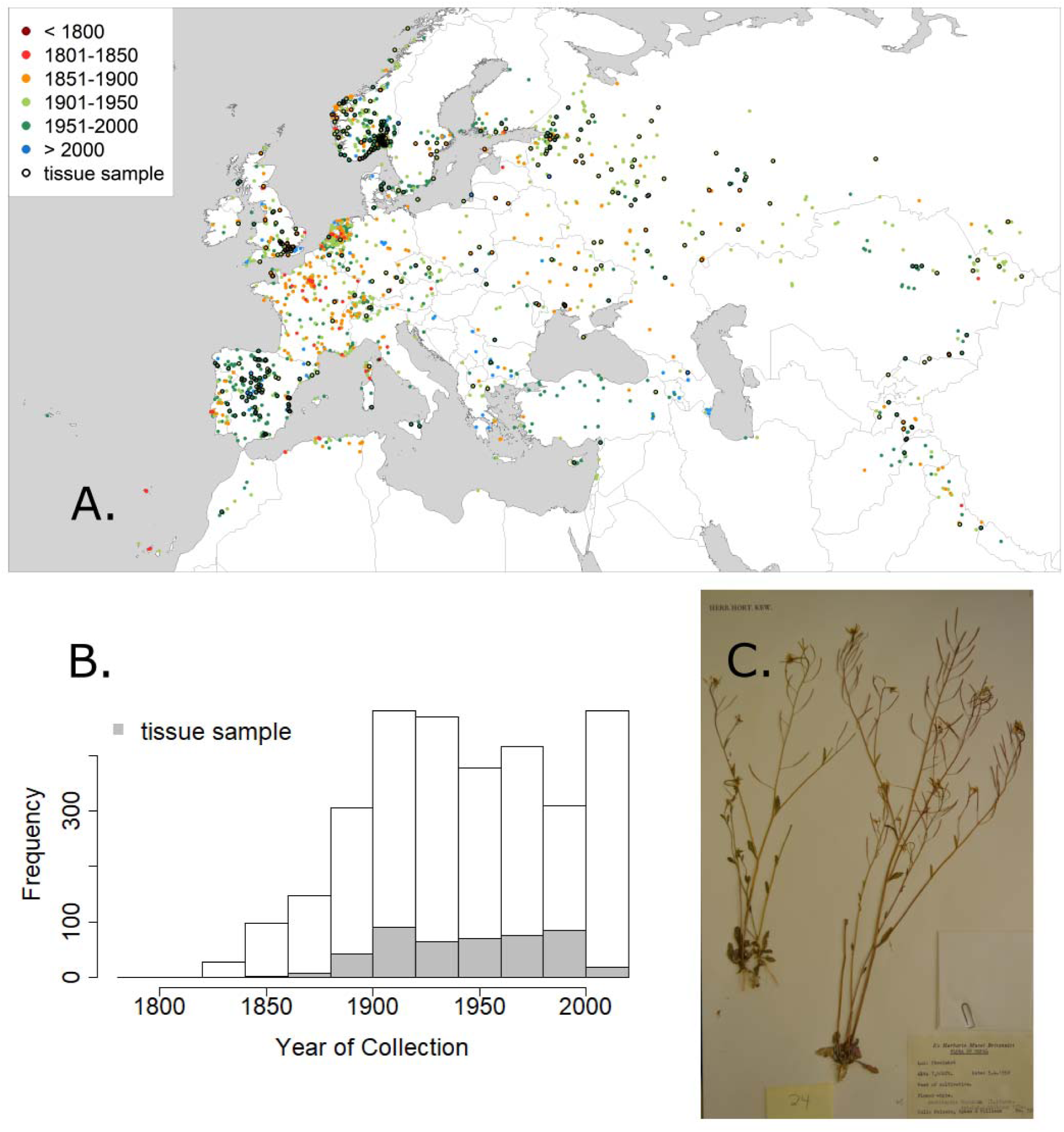
(A) Locations of collections used in our analysis. Color of circle corresponds to year of collection. Accessions that were sampled for tissue are outlined in black. (B) Distribution of years of collection. (C) Sample herbarium record from Nepal on April 5, 1952.

### Correlations among phenotypes (question 1)

We found generally weak correlations among phenotypes of *Arabidopsis* individuals (Question 1). The first two principal components explained only 36.2% and 24.1%, respectively, of the variance in the five phenotypes of Δ^13^C, δ^15^N, date of collection, C:N, and PTU (N = 397). The first principal component corresponded to a negative correlation between C:N versus day of collection (bivariate r = −0.189) and PTU (bivariate r = −0.101). Inspecting the relationship between collection date and C:N further revealed a triangle shape (Figure 2B). That is, there were no late-collected individuals with high C:N, potentially indicating that the late-collected plants that we hypothesize are spring/summer annuals also exhibit fast growth strategies. Plants with the lowest C:N have a less negative relationship to day of collection than plants with a higher C:N, and ANOVA showed there to be a significant difference between the slopes of the regression of the 25^th^ and 75^th^ percentiles of C:N (p = 0.0002, Figure S2) (Koenker & Koenker, 2011). When allowing the relationship between C:N and phenology to vary spatially (GAM with spatially varying coefficients), we found both date of collection and PTU were negatively correlated with C:N across the *Arabidopsis* native range, but this correlation was not significantly different from 0 (the 95% confidence interval included 0) (Figure S3). The second PC corresponded to a negative correlation between Δ^13^C and δ^15^N (bivariate r = −0.218). C:N and leaf proportion N are highly correlated (bivariate r = −0.816, Figure S1), so we focus on C:N. See supplementary material for leaf N results (Figure S16 and S17).

**Figure 2:**
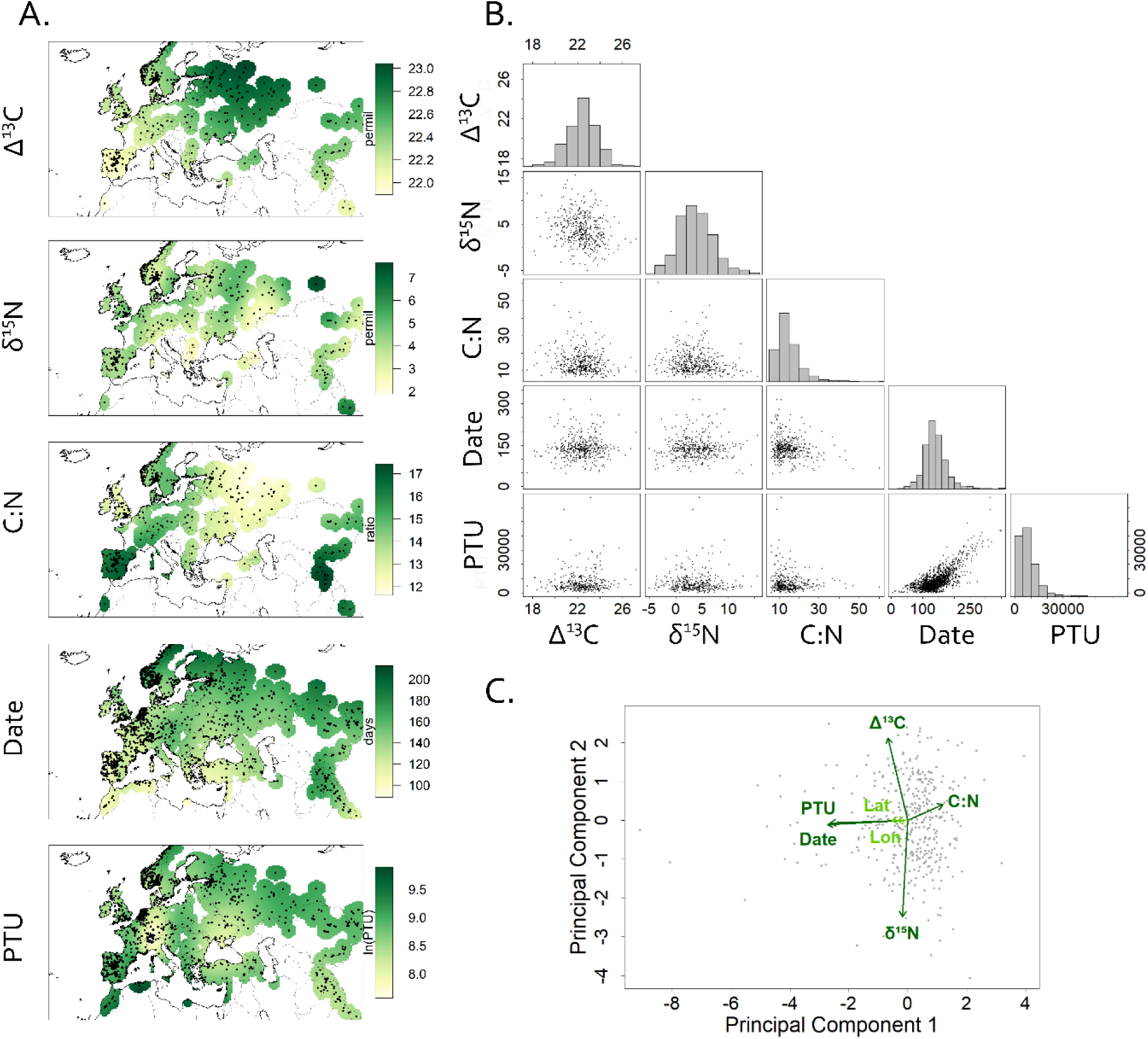
(A) Variation in phenotypes across the native range of *Arabidopsis* for Δ^13^C, δ^15^N, C:N, collection date, and photothermal units (PTU) at collection. Color indicates the fitted mean value (spatially-varying intercept) of the phenotype from the year-only model. For example, collection date is earlier in the Mediterranean in comparison to other regions; however, PTUs are lowest in the central part of the range. (B) Correlations between phenotypes in this study. Histograms of the measured values of each phenotype are plotted along the diagonal. (C) PCA of phenotypes. Correlations of phenotypes with principal components are plotted as arrows, with length multiplied by 3 for ease of viewing. Latitude (correlation with first PC r = −0.182) and longitude (correlation with first PC r = −0.115) are plotted for geographic context, though they were not included in PCA. Arrows for latitude and longitude are scaled equally to the arrows for phenotype correlations.

### Spatial variation in long-term average phenotypes (questions 2 and 3)

We visualized spatial diversity in phenotypes by plotting the spatial intercept surfaces in the year only models (Figure 2A). All phenotypes showed significant spatial variation (all GAM smooth terms significantly different from zero). Δ^13^C was lower in the Iberian Peninsula and higher in Russia (GAM smooth term, p = 0.0002). δ^15^N varied across the range, but with less pronounced spatial gradients (GAM smooth term, p = 0.01). C:N was higher in the Iberian Peninsula and central Asia and lower in Russia (GAM smooth term, p = 9e-05). Collection day was earlier along the Atlantic coast and Mediterranean (GAM smooth term, p = <2e-16). Despite this, PTU at collection still was higher in the Mediterranean region as well as at far northern, continental sites (GAM smooth term, p = <2e-16).

### Temporal change in phenotypes

Several phenotypes have changed significantly across large regions over the study period (1794-2010, Figure 3 above). For example, C:N ratio increased in later years in much of southwestern Europe. δ^15^N decreased significantly throughout most of the range. Collection date and PTUs became significantly later in many regions from the Mediterranean to Central Asia, although collection date became significantly earlier in the extreme south (Morocco and Himalayas). There was no significant temporal trend in Δ^13^C (not shown).

**Figure 3:**
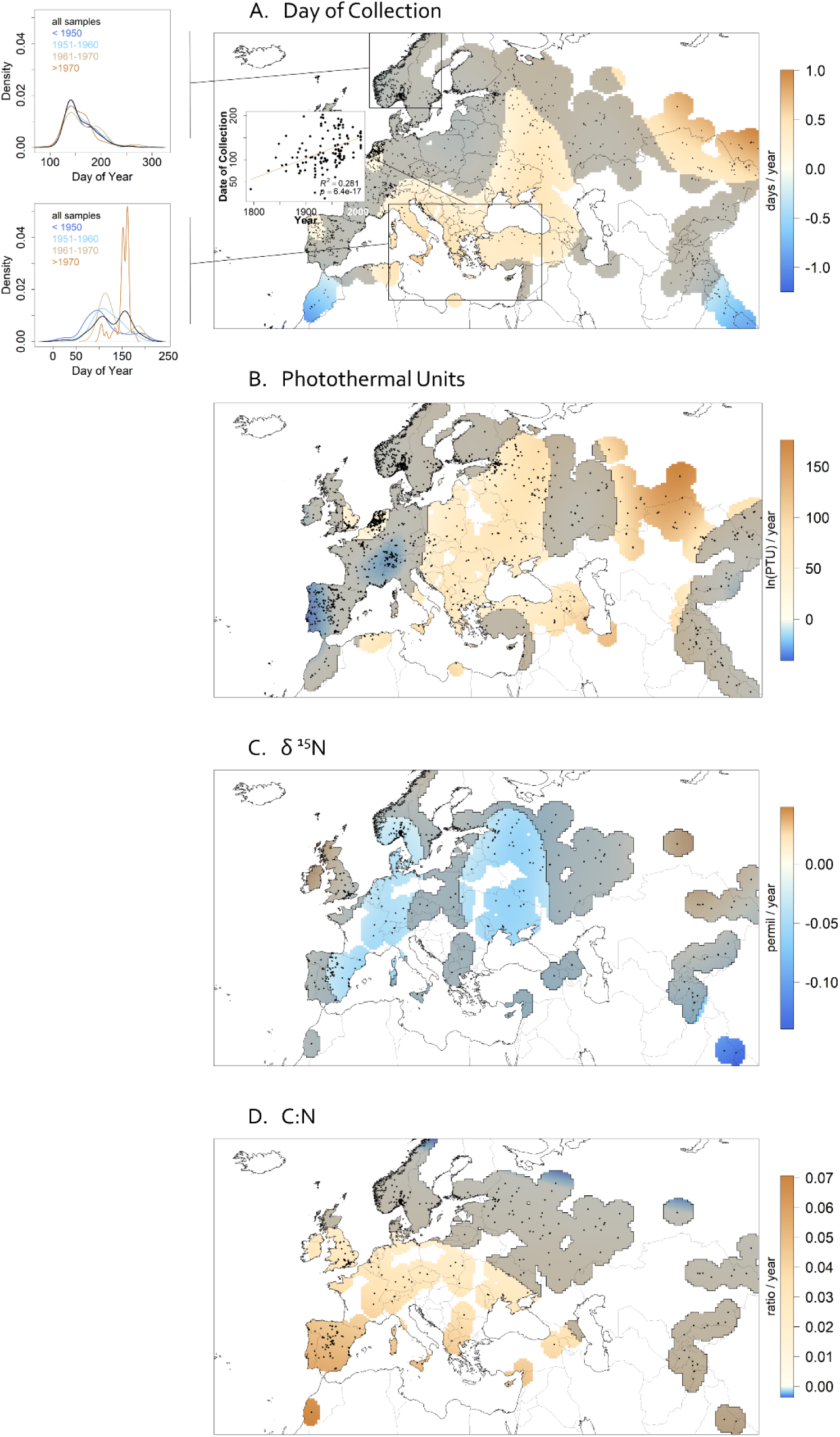
Change in phenotypes across years for collection date (A), photothermal units (B), δ nitrogen (C), and C:N (D). Color indicates the value of the coefficient for year in the model excluding climate variables, gray shading indicates regions where estimated coefficient is not significantly different from 0. For example, day of collection and photothermal units have significantly increased over time in most of the range, but with some exceptions for day of collection in the south. Inset scatterplot in A shows the significant increase in collection date with year for samples in the boxed Mediterranean region. Plots to the left of A show the density of collection dates through the year remains stable through time for Scandinavian collections within the boxed region (top) but shift toward more collections late in the year in the boxed Mediterranean collections (bottom).

The year trends in phenotypes across the study period were likely partly related to underlying climate variation. However, collection date, C:N, and δ^15^N were still significantly associated with year of collection even when accounting for temporal anomalies in climate from 1901-2010 (Figures S6, S12, S14). PTUs were even more negatively related to year of collection after controlling for temporal anomalies (Figure S8). A notable discrepancy is that Iberian collections were collected significantly earlier in later years when yearly climate anomalies were accounted for (Figure S6). There was still no significant temporal trend in Δ^13^C.

### Phenotype associations with spatiotemporal climate gradients (questions 2 and 3)

#### Date of collection

In years (temporal climate anomaly models) with a relatively warm April plants were collected significantly earlier (Figure 4A). Similarly, in locations (spatial climate models) with warmer temperatures plants were on average collected earlier in the year, though in many regions these coefficients were non-significant (Figure S7A, B). We also tested associations with July aridity index (precipitation/PET) and found that plants were collected significantly earlier in years (temporal climate anomaly models) with dry summers in Central/Eastern Europe (Figure 4B).

**Figure 4:**
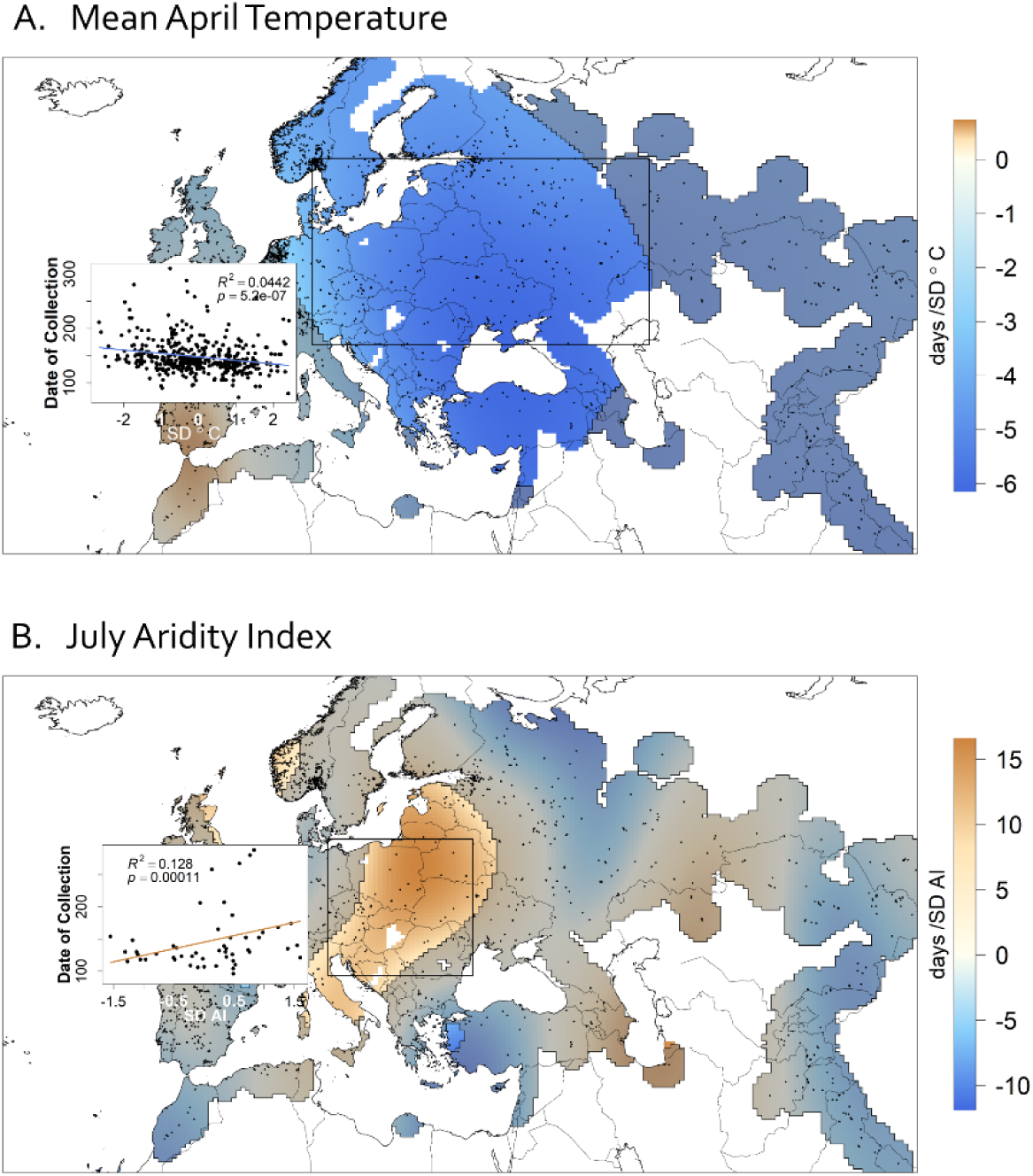
Association between collection day of *Arabidopsis* temporal mean April temperature anomalies (A) and July aridity index anomalies (B) (compared to 50-year average). Color indicates the value of the coefficient of the April mean temperature anomaly or July aridity index anomaly term. In years where April was warmer (positive anomalies), plants were collected earlier (a negative relationship). In wetter years, plants were collected later in Eastern Europe. Shading indicates regions where estimated coefficient is not significantly different from 0. Scatterplots of phenotype measures for individuals within the boxed areas show a decreasing collection date with mean April temperature anomaly and an increasing collection date with July aridity index anomaly in Eastern Europe and a decreasing collection date in Central Asia.

#### Photothermal units

To standardize spatiotemporal variation in developmental periods, we also modeled climate associations with PTUs. As expected, there were few areas where temperature anomalies were significantly associated with PTUs, likely due to the ability of PTUs to account for plastic responses (Figure S8). However, in some areas, accumulated PTUs at collection changed significantly in association with spatial temperature gradients, perhaps indicating spatial genetic differences in phenology (Figure S9). Locations with warmer Aprils had plants collected at more PTUs in East-Central Europe and fewer PTUs around the Aegean and Northern Asia. In the spatial climate model, plants from wetter areas in the Mediterranean were collected at lower PTUs (Figure S8, S9).

#### Δ ^13^Carbon

Wet summers were not significantly related to Δ^13^C in any region in either the temporal climate anomaly or spatial climate models (Figure S10, S11). Spatial variation in mean April temperatures was not significantly related to Δ^13^C, but plants from locations of colder Januaries did have lower Δ^13^C in Northern Asia and the Iberian Peninsula. (Figure S11B). Although elevation was not included in final models for reasons discussed above, replacing year with elevation in the temporal climate anomaly model showed a significant negative association between elevation and Δ^13^C (Figure S18D). Including elevation reduced the significance of the relationship between spatial variation in January temperature and Δ^13^C.

#### δ ^15^Nitrogen

δ^15^N was significantly higher in wetter years in Iberia, Asia, and Central Europe, (Figure S12C), but lower in the North of France. Spatial variation in minimum January temperatures was significantly positively related to δ^15^N around the North Sea (Figure S13B).

#### Leaf C:N

For the temporal climate anomaly model, plants collected in years with warmer winters in Iberia had significantly lower C:N ratios (Figure S14B. Leaf C:N differed in response to April mean temperature and January minimum temperature among locations (spatial climate models, Figure S15), although the patterns were mostly insignificant.

## Discussion

Widely distributed species often exhibit considerable phenotypic diversity, a large portion of which may be driven by adaptive plastic and evolutionary responses to environmental gradients. Previous studies of intraspecific trait variation in response to environment have tended to focus on genetic variation of environmental responses in common gardens (*e.g.* Wilczek *et al*. 2009; Kenney *et al*. 2014), temporal trends in phenology from well-monitored sites (*e.g.* CaraDonna et al., 2014), or field sampling of individuals from a small number of sites (*e.g.* Jung, Violle, Mondy, Hoffmann, & Muller, 2010). Here, we complement this literature by studying change in traits across an entire species range over two centuries, giving us a window into drivers of intraspecific diversity and regional differences in global change biology. From the accumulated effort contained in natural history collections, we tested hypotheses about variation in life history and physiology in response to environment. We observed modest evidence of coordinated phenological-physiological axes of variation for *Arabidopsis* in nature. We found later flowering times and higher accumulated photothermal units over the study period across most of the range and lower δ^15^N and higher C:N in more recent collections. Additionally, we observed distinct regional differences in phenology, Δ^13^C, and C:N in response to rainfall and temperature, potentially due to genetic differences among populations.

### Intraspecific variation in life history and physiology shows little coordination along a single major axis (question 1)

We found little evidence for tight coordination among studied phenotypes, fitting with some past studies that found weak to no support for a single major axis in intraspecific trait variation in response to environment across diverse plant growth forms (e.g. Albert *et al*. 2010; Wright & Sutton□Grier 2012). Common garden experiments often find substantial genetic covariation between the traits we studied possibly due to pleiotropy or selection maintaining correlated variation (Des Marais et al., 2012; Kenney et al., 2014; McKay et al., 2003). By contrast, the massively complex environmental variation organisms experience in the wild may combine with genotype-by-environment interactions to generate high dimensional trait variation among individuals in nature.

Nevertheless, we found modest evidence of a life history-physiology axis: plants collected later in the year had low leaf C:N, indicative of a fast-growing resource acquisitive strategy with low investment in C for structure and high investment in N for photosynthesis. The C:N/collection date axis is probably not due to later collections being at later developmental stages, since we would expect plants collected later in development to have allocated nitrogen away from leaves, lowering C:N. Instead, a strategy of higher leaf C:N may be adaptive for rapid-cycling plants germinating and flowering within a season (spring/summer annuals), which we expect to be collected later in the year due to later germination, contrasted with slower-growing genotypes known to require vernalization for early spring flowering over a winter annual habit. Indeed, Des Marais et al. (2012) found that vernalization-requiring (winter annual) *Arabidopsis* genotypes had lower leaf N than genotypes not requiring vernalization for flowering, the latter of which could also behave as spring or summer annuals.

The negative correlation we observed between Δ^13^C and δ^15^N has been reported by other authors and suggested to be a result of independent responses to multiple correlated environmental variables rather than a biological constraint. Environmental variables with opposing effects on Δ^13^C and δ^15^N include soil, temperature, and rainfall patterns (Hartman & Danin, 2010; Liu et al., 2007; Peri et al., 2012), due to depletion of soil N and changes in stomatal opening, and atmospheric carbon (Bloom, Burger, Asensio, & Cousins, 2010), which increases carbon uptake while suppressing nitrate assimilation.

### *Arabidopsis* life history and physiology vary across spatial environmental gradients, suggesting adaptive responses to long-term environmental conditions (question 2)

Geographic clines in traits in nature may be due to adaptive responses to environment. However, the spatial differences in traits or trait changes through time we observed are difficult to ascribe to genetic or plastic causes because of unknown genotype-environment interactions in the field and the confounding of environmental gradients and population genetic structure. The 1001 Genomes Project identified genetic clusters of *Arabidopsis* that were somewhat geographically structured but noted that these clusters overlapped and were distributed across a wide range of environments (Alonso-Blanco et al., 2016) (see Figure S5 for a map of the clusters). The patterns of significant phenotype-environment relationships we observed spanned multiple genetic clusters, making it unclear how much of a role these broad genetic groupings play in determining environmental response. The phenotype-environment relationships we observed followed our expectations for how phenology could affect fitness through earlier flowering times in response to warmth and both earlier and later flowering in response to drier environments. Physiological traits were less well aligned with our predictions for adaptive response; we did not find low Δ^13^C and high C:N associated with environments or years of water stress.

### Physiology, lack of correspondence to the Leaf Economic Spectrum

The Leaf Economic Spectrum and fast/slow life history predictions were not well supported by our results for how C:N, Δ^13^C, and δ^15^N respond to climate, since we saw both positive and negative trends with temperature and aridity depending on geographical region. This may be due to the intraspecific nature of our study, as opposed to the interspecific data often used to support the LES (Albert et al., 2010; Elmore, Craine, Nelson, & Guinn, 2017). In addition, our study may have overlooked the effects of edaphic conditions on C:N and δ^15^N. C:N over most of the native range was insignificantly related to spatial and temporal gradients of temperature and aridity index (July precipitation/PET) but increased with year as seen in grassland communities due to rising C_a_ (Gill et al., 2002). Likewise, δ^15^N over most of the range neither decreased with aridity index nor responded to temperature as expected, but did decrease with year as previously reported (McLauchlan, Ferguson, Wilson, Ocheltree, & Craine, 2010), possibly due to CO_2_ enrichment.

Similarly, we did not see strong relationships between aridity index and Δ^13^C. Δ^13^C was expected to be related to rainfall and temperature due to Δ^13^C being a proxy for stomatal gas exchange (Diefendorf et al., 2010; Farquhar et al., 1989). There are at least three potential explanations for weak Δ^13^C relationships with climate. First, we observed both positive and negative trends for aridity and date of collection, consistent with the hypothesis that *Arabidopsis* exhibits both drought escaping and drought avoiding genotypes. The phenological response to moisture of rapid flowering (drought escape strategy) could confine growth to periods of high moisture, obviating any stomatal closure in response to soil drying (and hence no effect on Δ^13^C). Stated simply, phenology and physiology cannot be treated as completely independent traits. Second, variation in plant traits we did not directly consider may affect Δ^13^C. Gas exchange and carbon assimilation depend in part on leaf architecture and physiology traits like venation, root allocation, and mesophyll conductance (Brodribb, Feild, & Jordan, 2007; Easlon et al., 2014; Schulze, Turner, Nicolle, & Schumacher, 2006), which could limit responses in Δ^13^C. For example, given the role of roots in sensing drought and triggering stomatal response (Christmann, Hoffmann, Teplova, Grill, & Muller, 2004), greater investment in roots could allow plants in relatively drier conditions to maintain open stomata, preventing decreases in C_i_ and leading to no observed climate effect on Δ^13^C. Third, elevated atmospheric partial CO_2_ could mitigate climate effects on Δ^13^C by increasing the efficiency of stomatal gas exchange (Drake, Hanson, Lowrey, & Sharp, 2017). Local investigations of the patterns we found could complement our results by characterizing the underlying ecophysiological and life history mechanisms driving intraspecific variation.

### Phenology, high variation across space

We found strong spatial gradients in two measures of phenology, suggesting that adaptive responses to climate drive long-term trait differences among regions. Locations that were warmer than average in either April or January corresponded to significantly earlier collection dates, consistent with temperature’s positive effect on growth rate (Wilczek et al., 2009). In addition, our models provided support that some phenological variation did reflect seasonality of moisture availability. We found that *Arabidopsis* was collected significantly earlier in years with dry summers in central Europe and at significantly lower PTU in regions of wet summers around the Mediterranean, suggesting drought escape or avoidance strategies, respectively, could be important in those regions. Alternatively, later collections in wetter years could be the result of multiple successful generations due to the extra rainfall. This ambiguity illustrates an important caveat in using collection date and PTU to study *Arabidopsis* in the wild. Because we cannot differentiate changes in total life span from changes in germination start date, shifts in phenology (the timing of an organism’s life to seasonal conditions) does not necessarily indicate shifts in life history. Nevertheless, the observed regional variation in the response of collection date and PTU to climate variables provides valuable insight into how different populations experience their environment and suggest areas for more direct study.

### Changes in *Arabidopsis* life history and physiology over the last two centuries track climate, suggesting adaptive responses (question 3)

Increasing global temperatures were expected to increase relative growth rate and hasten germination, decreasing flowering time as measured by collection date. In addition, atmospheric CO_2_ enrichment was expected to increase Δ^13^C (Drake et al., 2017) and C:N and decrease δ^15^N (Bloom et al., 2010). With increasing environmental nitrogen due to human activity (Galloway et al., 2004), we expected δ^15^N to decrease.

Our findings were largely consistent with these hypotheses in the year models for leaf physiology. Δ^13^C did not significantly change through time across the native range, which could be due to life history shifts as mentioned above or differential response to aridity gradients across the locations sampled masking the effect of elevated CO_2_ (Drake et al., 2017). Geographic variation in the strength of the relationships for other traits could be due to underlying genetic variation or interaction with environmental factors we did not account for. Nevertheless, C:N increased and δ^15^N decreased as expected across large portions of the native range.

For collection date and PTU, however, our models returned the surprising result of later collection rather than earlier, despite earlier collections in warmer years. The fact that the relationship between warmth and collection date was spatially variable, and insignificant in some regions, may indicate areas of contrasting phenological response, perhaps due to lost vernalization signal or variable effects on germination (Burghardt, Edwards, & Donohue, 2016). Variation across space in phenological response to climate change has been shown before and may be due to genetic differences among populations or due to interactions with other environmental variables (Park et al., 2018). *Arabidopsis* is known to complete a generation within a single season, climate permitting, and warmer climates may even allow for fall flowering (Fournier-Level et al., 2013; Wilczek et al., 2009). If warmer temperatures enable a greater number of spring or summer germinants to flower before winter in regions such as Central Europe, we would expect to see later collection dates in more recent years (Burghardt et al., 2015). Regions that did not show later collection dates through time might be limited in generational cycles due to summer drought or very short growing seasons. For instance, early flowering in the spring has been implicated as an important strategy for *Arabidopsis* in the Iberian Peninsula to escape seasonal heat and water limitation that curtail growth in later months (Wolfe & Tonsor, 2014).

Our findings of later collection dates through the study period (1798-2010) may surprise some readers due to previously observed acceleration of temperate spring phenology (Parmesan & Yohe, 2003). However, we modeled changes in mean phenological response to environment, which can be weakly related to either tail of phenology trait distributions (CaraDonna et al., 2014). Individuals on the extremes, such as first-flowering individuals, are often the primary focus of studies showing accelerated spring phenology in recent years. Why might *Arabidopsis* flower later even as global temperatures rise? First, anthropogenic land use change may drive phenology by favoring spring germinants, *e.g.* if disturbances favor faster life cycles. Second, warming climate or increasing atmospheric pCO_2_ may favor alternate life histories by increasing relative growth rate, thus allowing spring or fall germinants to complete their life cycle before conditions degrade at the end of a growing season. Later collections in more recent years might represent an increasing proportion of fast-growing spring or summer annuals as opposed to winter annuals. Whatever the cause of *Arabidopsis* flowering later, these phenological changes may have important ecological effects, such as altered biotic interactions.

We found evidence that trait correlations may be changing through time. For example, in some regions both C:N and date of collection have significantly increased over the past 200 years (around the Eastern Mediterranean), while in other regions date of collection decreased while C:N increased over time (Morocco and the Iberian Peninsula) (Figure 1). If the negative relationship between leaf C:N and flowering time that we observed is truly an axis of adaptive tradeoff between fast and slow life histories, this tradeoff may be changing at different rates among regions with time. Changing environments might reshape biological constraints on adaptive plant responses (Sgrò & Hoffmann, 2004).

### Our approach, technical limitations in herbaria data to surmount in future studies

Understanding how environmental variation drives the intraspecific diversity in broadly distributed species has been challenging due to logistics of large spatiotemporal scales. However, advances in digitization of museum specimens and the generation of global gridded spatiotemporal environmental data are opening a new window into large scale patterns of biodiversity. One challenge of herbarium specimens is that they typically present a single observation of a mature, reproductive individuals. Thus, these specimens contain limited information on phenology and physiology at earlier life stages (*e.g.* seedling plants), which can have subsequently strong impacts on later observed stages. Use of developmental models (Burghardt et al., 2015) might allow one to backcast potential developmental trajectories using herbarium specimens and climate data, to make predictions about phenology of germination and transition to flowering. In addition, herbaria collections are often biased by factors such as geography, species, and climate (Daru et al., 2018; Loiselle et al., 2008). Hierarchical sampling through repeated collections in the same region could improve the confidence of our model in representing phenotypic change through time.

Generalized additive models are a flexible approach to model phenotype responses to environment that might differ spatially among populations (MacGillivray et al., 2010). These models allow the data to inform on spatial variation in the trends studied, unlike approaches that bin individuals into discrete and arbitrarily bounded regions. Herbarium records represent imperfect and biased samples of natural populations (Daru et al., 2018), and future efforts may benefit from additional information that might allow us to account for these biases. Here, we sampled a very large number of specimens across continents and centuries, perhaps reducing the effect of biases associated with specific collectors. Nevertheless, as museum informatics advance it may become possible to explicitly model potential sources of bias, for example those arising from collecting behavior of specific researchers.

### Conclusion

Widely distributed species often harbor extensive intraspecific trait diversity. Natural history collections offer a window into this diversity and in particular allow investigation of long-term responses to anthropogenic change across species ranges. Here we show that spatiotemporal climate gradients explain much of this diversity but nevertheless much of the phenotypic diversity in nature for the model plant remains to be explained.

## Acknowledgements

Hundreds of professional and amateur botanists collected the specimens studied here. The staff at herbaria at Oslo Natural History Museum, Kew Gardens, Real Jardin Botanico, Komarov Botanical Garden, the New York Botanical Garden, and the British Museum of Natural History gave permission for tissue sampling. Michelle Brown provided essential assistance in collecting herbarium tissue. Jason Bonnette helped coordinate sample preparation. Major assistance in digitizing of specimens was provided by Patrick Herné. Eugene Shakirov aided in translating Russian specimen labels. Data from the MNHN in Paris were obtained thanks to the participatory science program Les Herbonautes” (MNHN/Tela Botanica) which is part of Infrastructure Nationale e-RECOLNAT: ANR-11-INBS-0004. Additional volunteers from the Atlas of Living Australia helped with digitization and georeferencing. Funding was provided by an Earth Institute fellowship to JRL.

The authors report no commercial or other relationships relevant to the content of this article that would represent a conflict of interest.

## Comparison of results to hypotheses

**Table 2:**
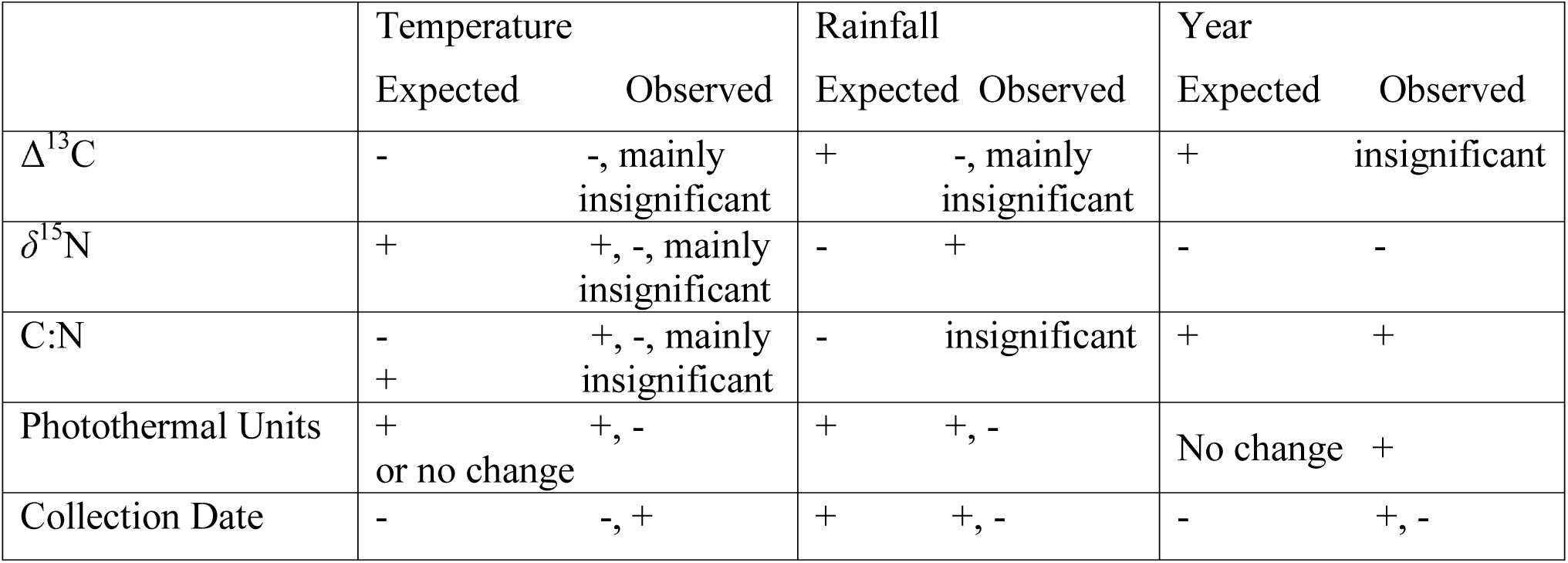
Expected phenotype responses to increases in temperature, rainfall, or year and observed model output. For some phenotypes a single trend over time was observed; however, most phenotypes showed variation in responses to temperature and rainfall across the range.

## Notes

#### Summary of Updates

Models were rerun with an updated version of climate data. Plots and text have been updated to reflect the results of the analysis.

