## Supplemental Material for "Effects of two centuries of global environmental variation on phenology and physiology of *Arabidopsis thaliana*"

**Supplement**

*Dataset*

Supplementary Table 1: Filters applied to Arabidopsis accessions, and final numbers of accessions considered for each phenotype.

| 3443 | Total instances of Arabidopsis with collection date* |
| --- | --- |
| 72 | Unable to visually check record |
| 41 | Removed due to collection date in midwinter (suspected unreliable) |
| 11 | Outside native range |
| 10 | Misidentified species, mostly *Arabis* spp. |
| 60 | Only fruits present or senesced plants |
| 118 | Only flowers present |
| 21 | Neither flowers nor fruits present |
| **3110** | Confident occurrences of Arabidopsis flowering/fruiting date |

Of these:

| 468 | Sent to Stable Isotope Facility |
| --- | --- |
| 2 | Included non-vegetative tissue in chemical analysis |
| 453 | δ^15^N, Δ^13^C, C:N, Proportion N |
| 1 | δ^15^N, Δ^13^C |
| 5 | δ^15^N, C:N, Proportion N |
| **459** | Samples with chemical phenotypes |

*There are 2 samples included that do not have a specific date of collection but were sampled for leaf chemistry

*Relationships between phenotypes*

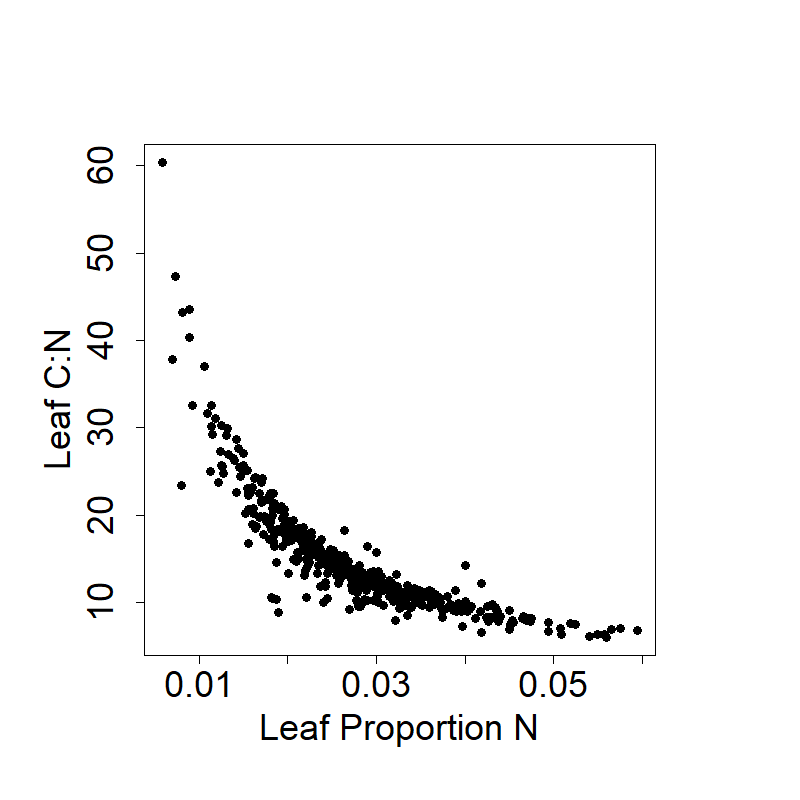

Figure S1: Proportion leaf nitrogen and leaf C:N ratio of the accessions sampled in our study show a strong negative association.

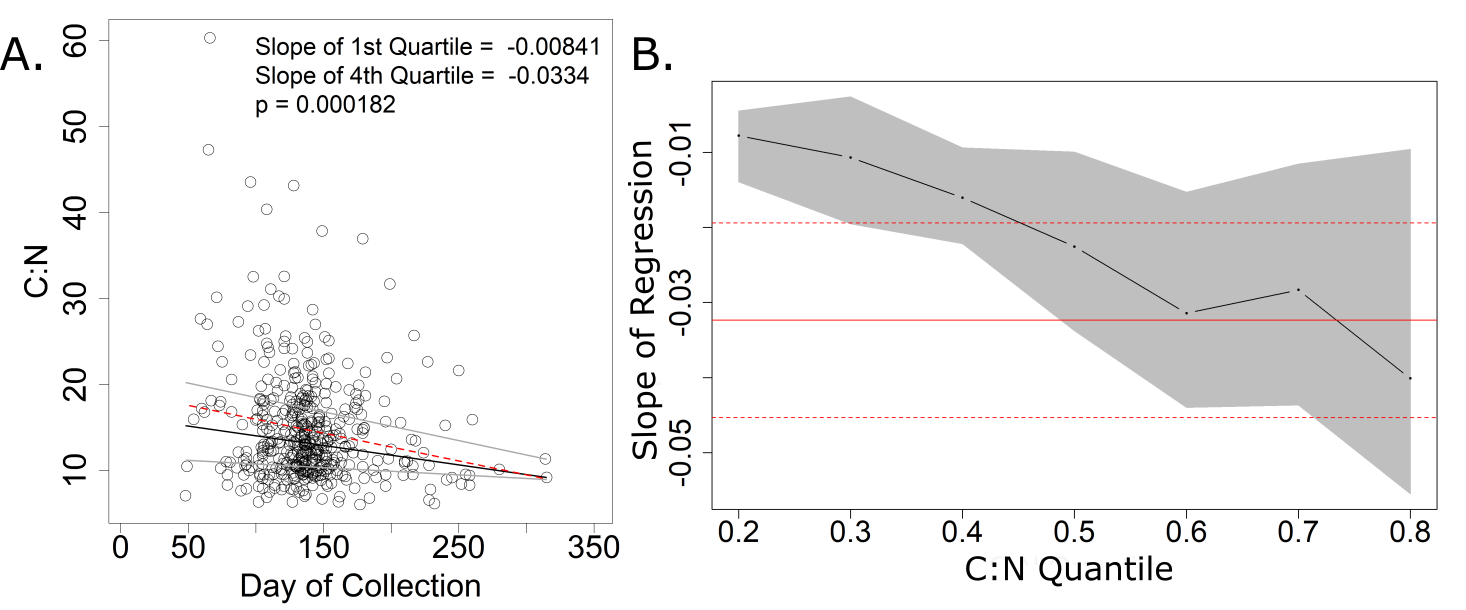

Figure S2: Quantile regression for C:N predicted by day of collection. No late collected accessions have a high C:N ratio. A) Gray lines indicate the regression of accessions in the 1^st^ and 4^th^ quartiles for C:N, while the black line indicates the regression of all data. The red line indicates the ordinary least squares line. ANOVA of the regressions of the 1^st^ and 4^th^ quartiles showed them to be statistically significantly different (p = 0.00018), suggesting that the highest C:N values respond more negatively as day of collection increases than lower C:N values. B) Slopes for accessions between the 20^th^ and 30^th^ percentile for C:N differ from the least squares estimate of all data, although the solution is non-unique. The red solid line shows the least squares estimate, while the dotted lines show the 95% confidence intervals. Black dots represent the calculated slope for each quantile of the C:N values.

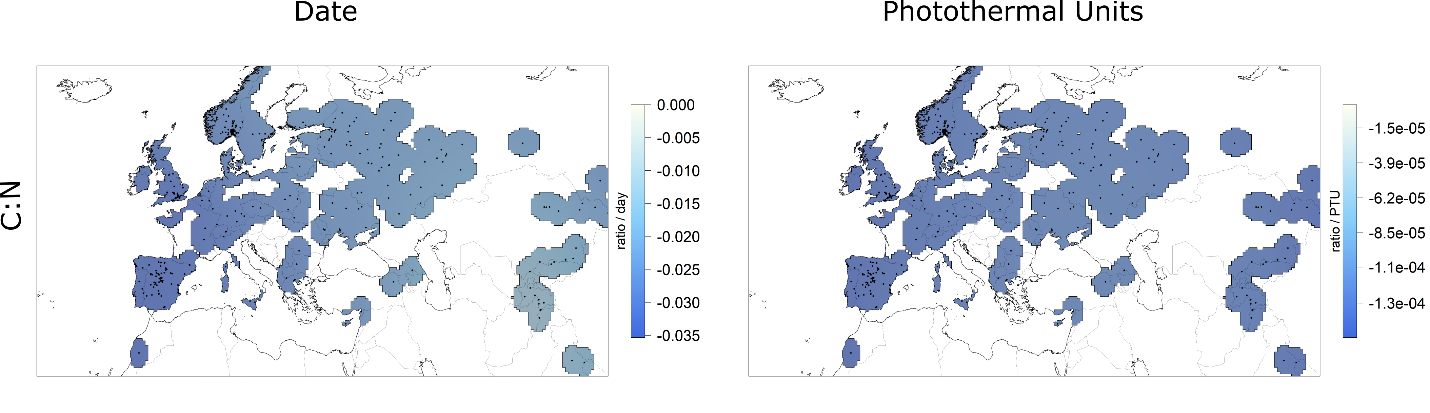
Figure S3: Relationship between selected phenotypes. C:N (dependent variable) may be negatively related to days and PTU at collection (independent variables) across the native range, but the linear regression slope does not differ significantly from 0 across the 95% confidence interval.

Model statistics:

C:N/Date

Parametric coefficients:

Estimate	Std. Error	t value	Pr(>|t|)

(Intercept)	18.515	1.162	15.93	<2e-16

Approximate significance of smooth terms:

edf	Ref.df	F	p-value

s(lon,lat):intercept	10.534	29	1.499	1.73e-12

s(lon,lat):dayofyear	2.931	29	0.505	0.000192

R-sq.(adj) =	0.16	Deviance explained = 18.5%

GCV = 35.409 Scale est. = 34.29	n = 458

AIC = 2934.946

C:N/PTU

Parametric coefficients:

Estimate	Std. Error	t value	Pr(>|t|)

(Intercept)	15.9193	0.4301	37.01	<2e-16

Approximate significance of smooth terms:

edf	Ref.df	F	p-value

s(lon,lat):intercept	11.784	29	2.347	8.18e-12

s(lon,lat):ptu	1.004	30	0.335	0.000853

R-sq.(adj) = 0.153	Deviance explained = 18.1%

GCV = 38.983 Scale est. = 37.635	n = 399

AIC = 2595.407

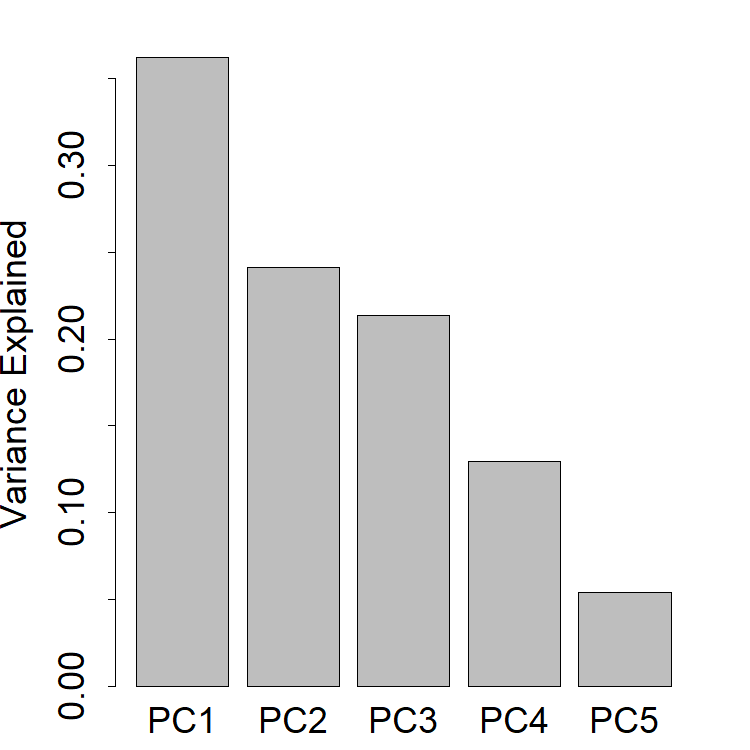

Figure S4: Proportion of variance explained by a Principal Components Analysis of five phenotypes in 397 samples.

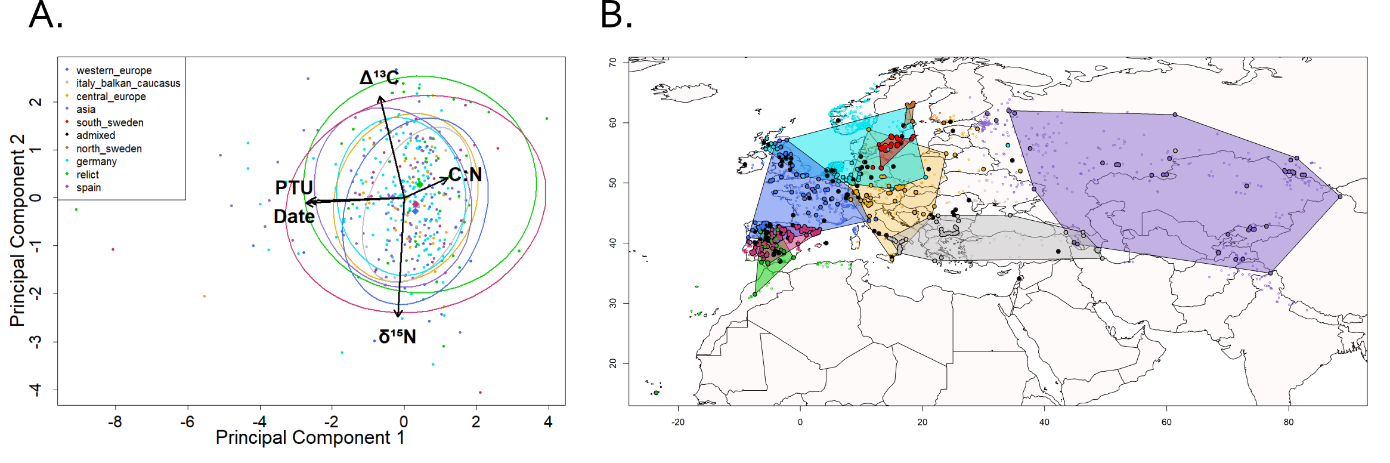
Figure S5:

(A) Herbarium samples assigned to the nearest geographic genetic cluster from the 1001 genomes project did not show strong patterns of genetic or geographic determination of phenotypic values in the principal component space. Ellipses include 75% of group members. (B) The genetic clusters suggested by the 1001 Genomes Project analysis. Large colored circles represent the locations of ecotypes sequenced by the 1001 Genomes Project and their group asisgnment, and small colored points indicate the accessions in our studu and their group assignment.

Polygon distributions for genetic clusters were created using adehabitat^1^ on groupings downloaded from the 1001 Genomes Project^2^. Accessions used in our analysis were assigned to the nearest cluster polygon. In the case of overlapping polygons, clusters were assigned in numeric order, causing no herbarium samples to be assigned to the South Sweden genetic cluster in this analysis. All African accessions were assigned to the relict group. Principal Component Analysis was performed as described in methods. The function dataEllipse from the package car^3^ was used to draw ellipses around 75% of group members for each cluster.

^1^Calenge, C. (2006) The package adehabitat for the R software: a tool for the analysis of space and habitat use by animals. Ecological Modelling, 197, 516-519.

^2^http://1001genomes.github.io/admixture-map/

^3^John Fox and Sanford Weisberg (2011). An {R} Companion to Applied Regression, Second Edition. Thousand Oaks CA: Sage. URL: http://socserv.socsci.mcmaster.ca/jfox/Books/Companion

*Additional maps of GAM parameter estimates*

How to interpret these plots: Each map shows the value of the slope of a covariate across space. In areas that are more blue, the covariate (e.g. April Mean Temperature, Year) is negatively related to the phenotype (e.g. Date of Collection). In areas that are more brown, the covariate is positively related to the phenotype. In areas where the 95% confidence interval of a slope included zero, we shaded the map in gray to indicate that the trend is not statistically significanly different from zero. For temporal anomaly models, where we used local climate anomalies, a positive value in the covariate indicated a higher value than the local average for that climate variable. Thus, areas that are brown show regions where higher than local average climate values in a year lead to an increase in the modelled trait and areas that are blue are regions where higher than average climate values lead to a decrease in the modelled trait. Locations of accessions used in fitting each model are plotted as black points on the map. For the reported model statistics, the following abbreviations are used:

tmn.1ano	standardized anomaly of January minimum temperature

tmp.4ano	standardized anomaly of April mean temperature

ai.7ano	standardized anomaly of July Aridity Index

ltmn2	normalized, 50-year average of January minimum temperature

ltmp2	normalized, 50-year average of April mean temperature

lai2	normalized, 50-year average of July Aridity Index

year2	normalized year of collection

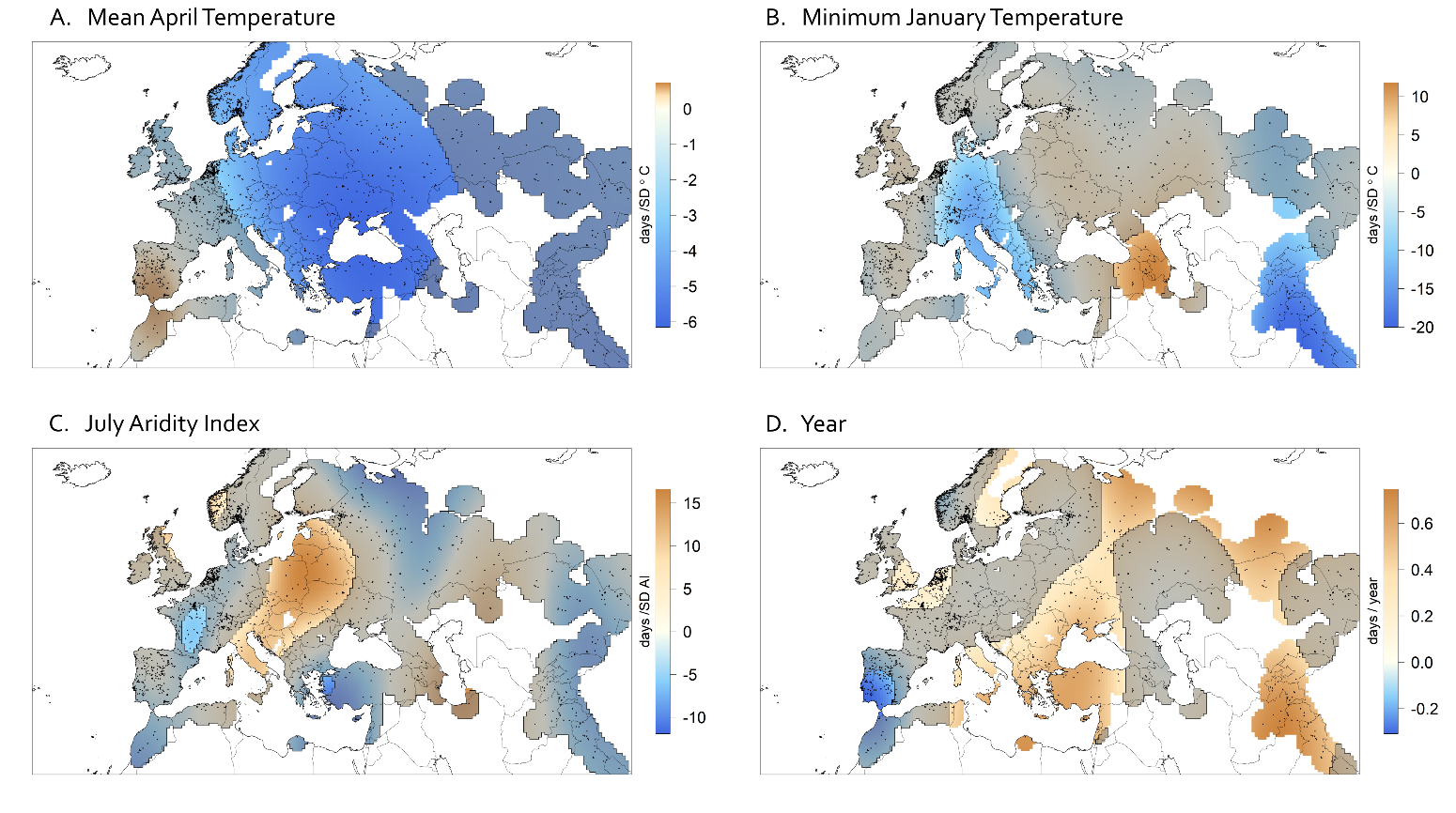
*Phenology temporal anomaly model*

Figure S6: Coefficient surfaces of mean April temperature (A), minimum January temperature (B), July Aridity Index (C), and year (D) in a temporal anomaly model of day of collection with a spatially varying intercept. For each climate variable, conditions in the year of collection are scaled to long term averages at that location. Shading represents areas where the 95% confidence interval of the estimated coefficent includes 0. Plants were collected earlier in years where spring was warmer. Warmer winters saw earlier collections in Central Europe and Asia but later collections in the Middle East. Rainfall was associated with later collections in Eastern Europe. In most of the range, plants are being collected later in later years, but in the Iberian Peninsula this trend is reversed.

Parametric coefficients:

Estimate	Std. Error	t value	Pr(>|t|)

(Intercept)	139.9964	0.8591	163	<2e-16

Approximate significance of smooth terms:

edf	Ref.df	F	p-value

s(lon,lat):intercept	26.02	29	18.771	< 2e-16

s(lon,lat):tmn.1ano	17.80	30	1.944	1.46e-07

s(lon,lat):tmp.4ano	4.71	30	0.823	7.79e-06

s(lon,lat):ai.7ano	21.95	30	1.464	0.00113

s(lon,lat):year2	20.43	30	3.098	1.10e-13

R-sq.(adj) = 0.265	Deviance explained = 29.1%

GCV = 1094.4 Scale est. = 1054.1	n = 2495

AIC = 24538.97

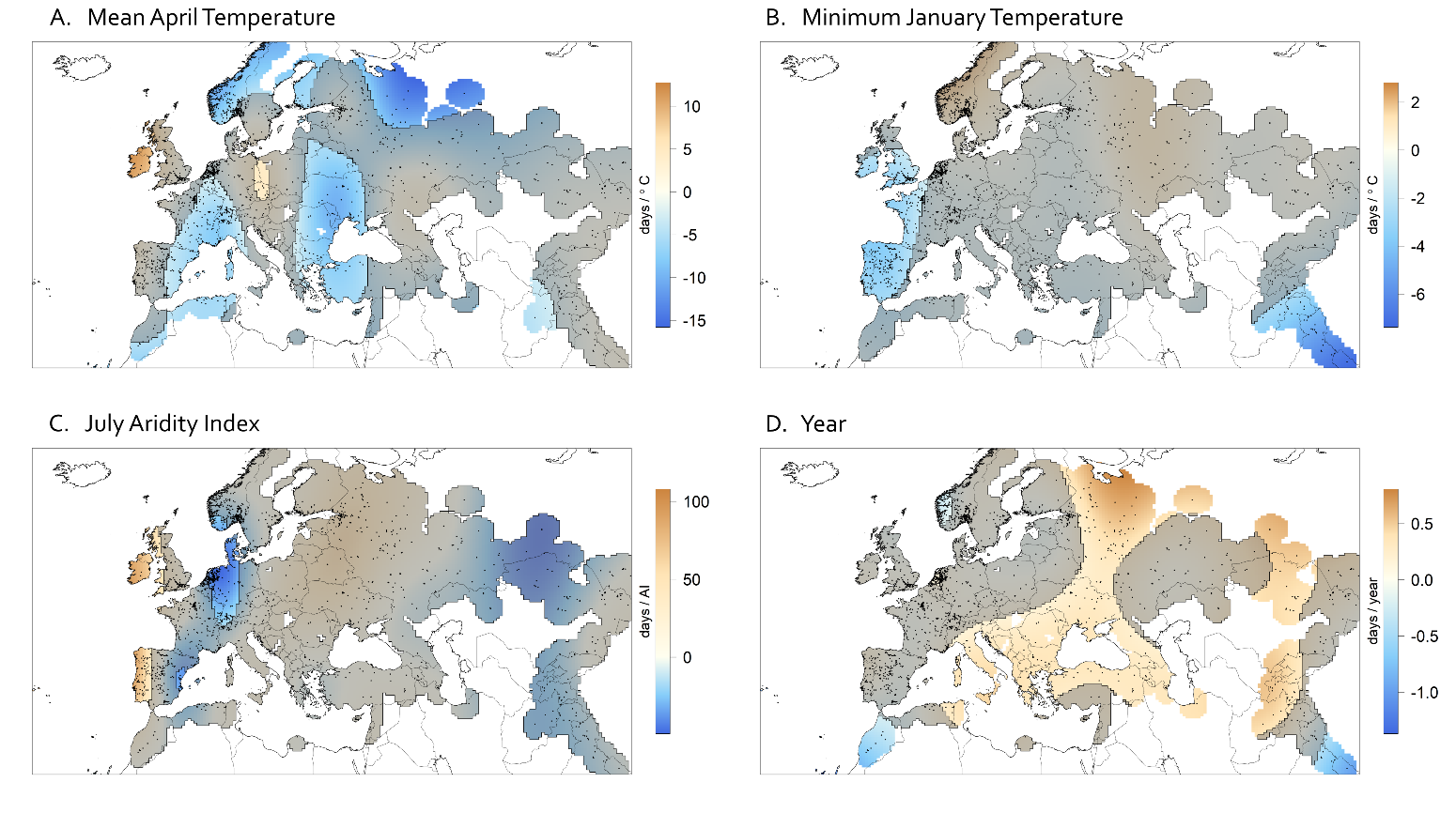
*Phenology spatial model*

Figure S7: Coefficient surfaces of mean April temperature (A), minimum January temperature (B), July Aridity Index (C), and year (D) in a spatial model of collection date with a non-spatially varying intercept. For each climate variable, 50-year averages at each location are used. Shading represents areas where the 95% confidence interval of the estimated coefficent includes 0. Plants are collected earlier in areas with warmer springs in much of the range. Across most of the range, plants are collected later in more recent years.

Parametric coefficients:

Estimate	Std. Error	t value	Pr(>|t|)

(Intercept)	143.775	2.197	65.43	<2e-16

Approximate significance of smooth terms:

edf	Ref.df	F	p-value

s(lon,lat):ltmn2	14.38	30	1.308	5.03e-06

s(lon,lat):ltmp2	25.12	30	4.253	< 2e-16

s(lon,lat):lai2	19.98	30	1.631	6.40e-06

s(lon,lat):year2	25.67	30	5.380	< 2e-16

R-sq.(adj) = 0.335	Deviance explained = 35.4%

GCV = 1054.5 Scale est. = 1025.3	n = 3108

AIC = 30453.99

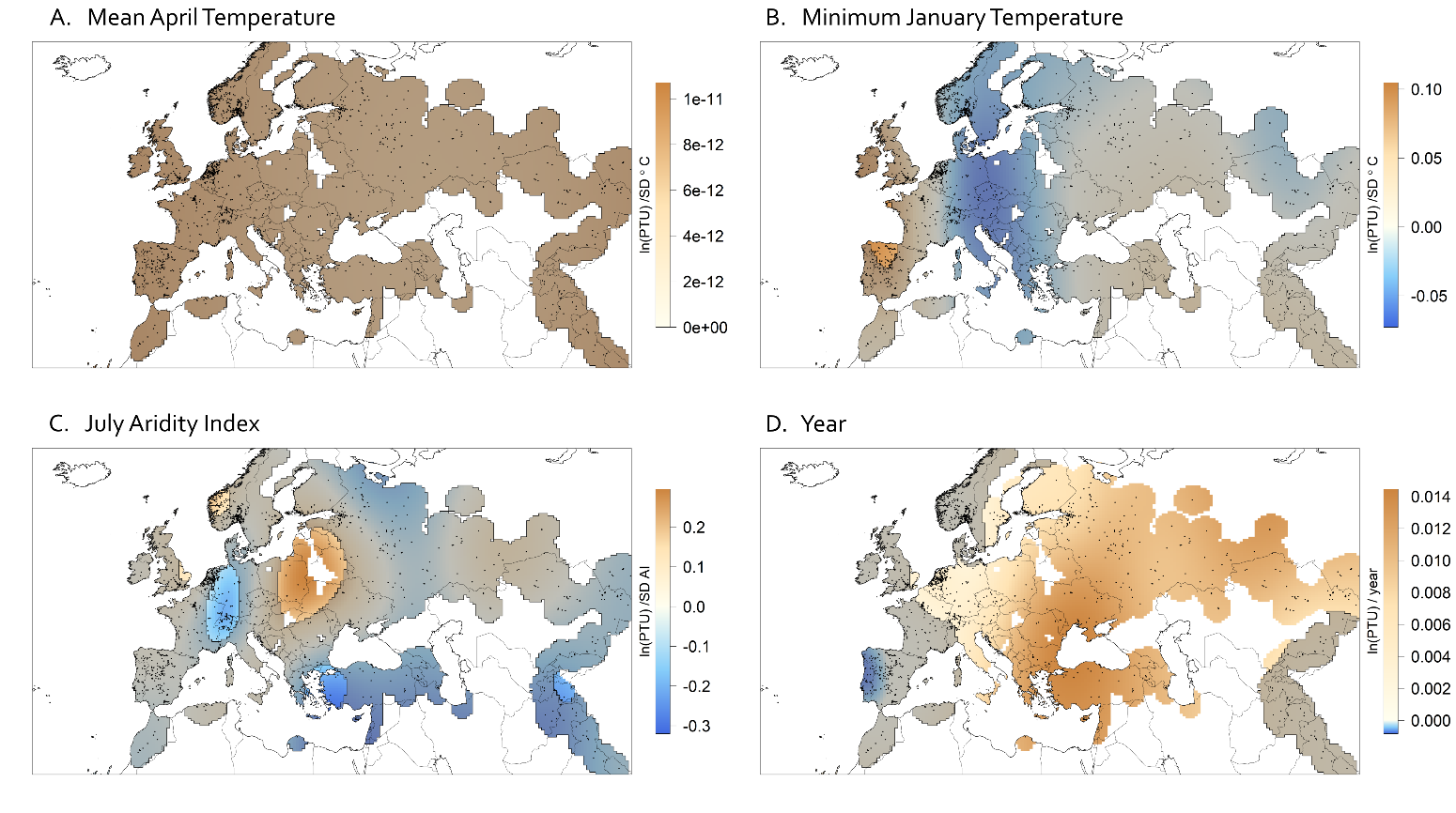
*Photothermal Units temporal anomaly model*

Figure S8: Coefficient surfaces of mean April temperature (A), minimum January temperature (B), July Aridity Index (C), and year (D) in a temporal anomaly model of Photothermal Units with a spatially varying intercept. For each climate variable, conditions in the year of collection are scaled to long term averages at that location. Shading represents areas where the 95% confidence interval of the estimated coefficent includes 0. In Eastern Europe, wetter years were associated with a greater accumulation of PTUs by collection, although the opposite was observed in West-Central Europe. Photothermal Units at time of collection increased through time across most of the range.

Parametric coefficients:

Estimate	Std. Error	t value	Pr(>|t|)

(Intercept)	8.67723	0.02005	432.7	<2e-16

Approximate significance of smooth terms:

edf	Ref.df	F	p-value

s(lon,lat):intercept	2.441e+01	29	11.476	< 2e-16

s(lon,lat):tmn.1ano	6.967e+00	29	0.386	0.079907

s(lon,lat):tmp.4ano	1.803e-09	30	0.000	0.784226

s(lon,lat):ai.7ano	1.912e+01	30	1.341	0.000918

s(lon,lat):year2	1.126e+01	30	3.368	< 2e-16

R-sq.(adj) = 0.194	Deviance explained = 21.4%

GCV = 0.66434 Scale est. = 0.64758	n = 2488

AIC = 6043.514

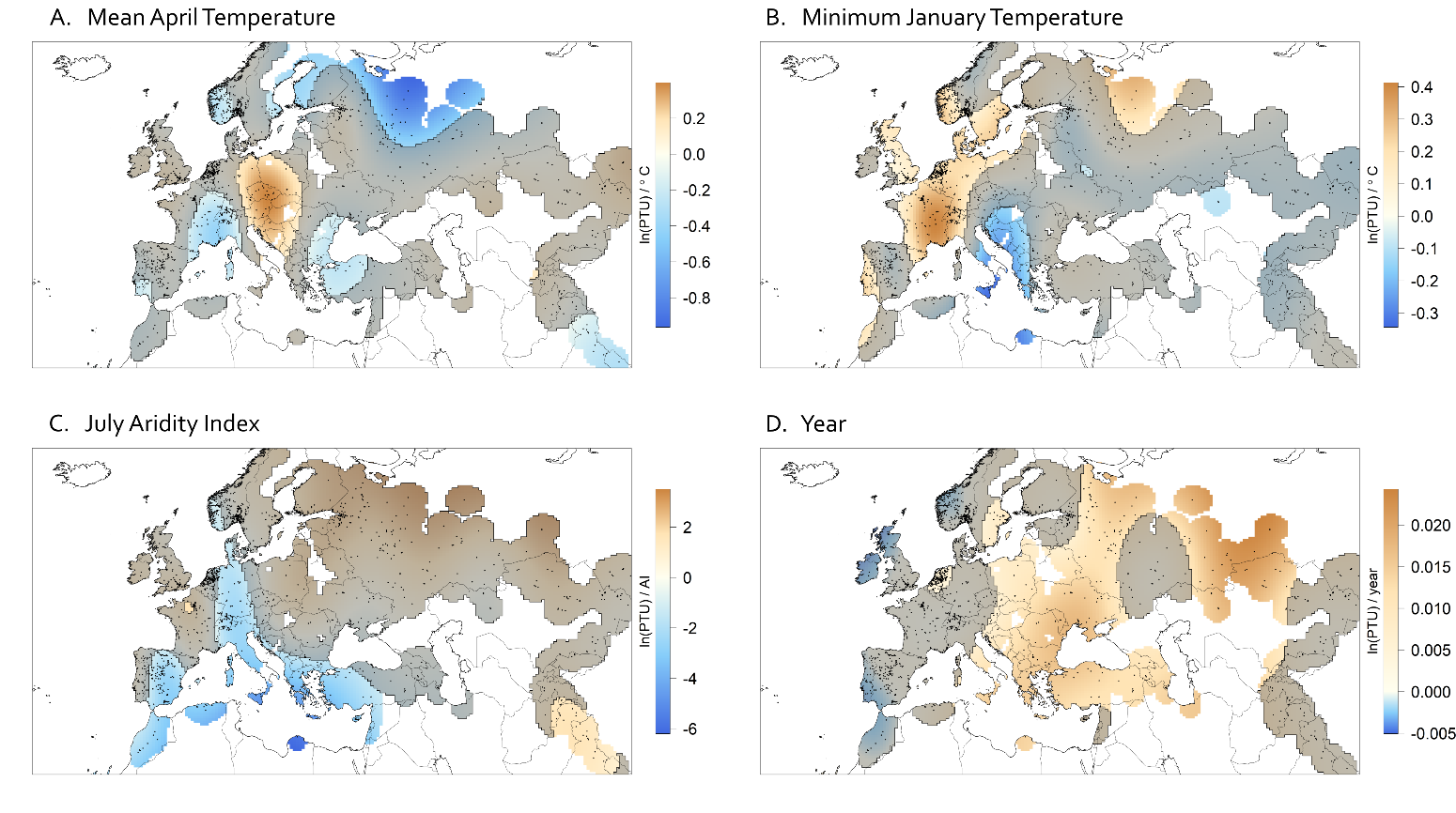
*Photothermal Units spatial model*

Figure S9: Coefficient surfaces of mean April temperature (A), minimum January temperature (B), July Aridity Index (C), and year (D) in a spatial model of Photothermal Units with a non-spatially varying intercept. For each climate variable, 50-year averages at each location are used. Shading represents areas where the 95% confidence interval of the estimated coefficent includes 0. The relationship with Photothermal Units and each climate variable is heterogenous through space. Photothermal Units at time of collection has increased in more recent years.

Parametric coefficients:

Estimate	Std. Error	t value	Pr(>|t|)

(Intercept)	8.22814	0.07245	113.6	<2e-16

Approximate significance of smooth terms:

edf	Ref.df	F	p-value

s(lon,lat):ltmn2	28.36	30	4.356	<2e-16

s(lon,lat):ltmp2	25.22	30	4.554	<2e-16

s(lon,lat):lai2	23.88	30	10.482	<2e-16

s(lon,lat):year2	21.52	30	4.987	<2e-16

R-sq.(adj) = 0.241	Deviance explained = 27.1%

GCV = 0.63576 Scale est. = 0.61018	n = 2485

AIC = 5924.457

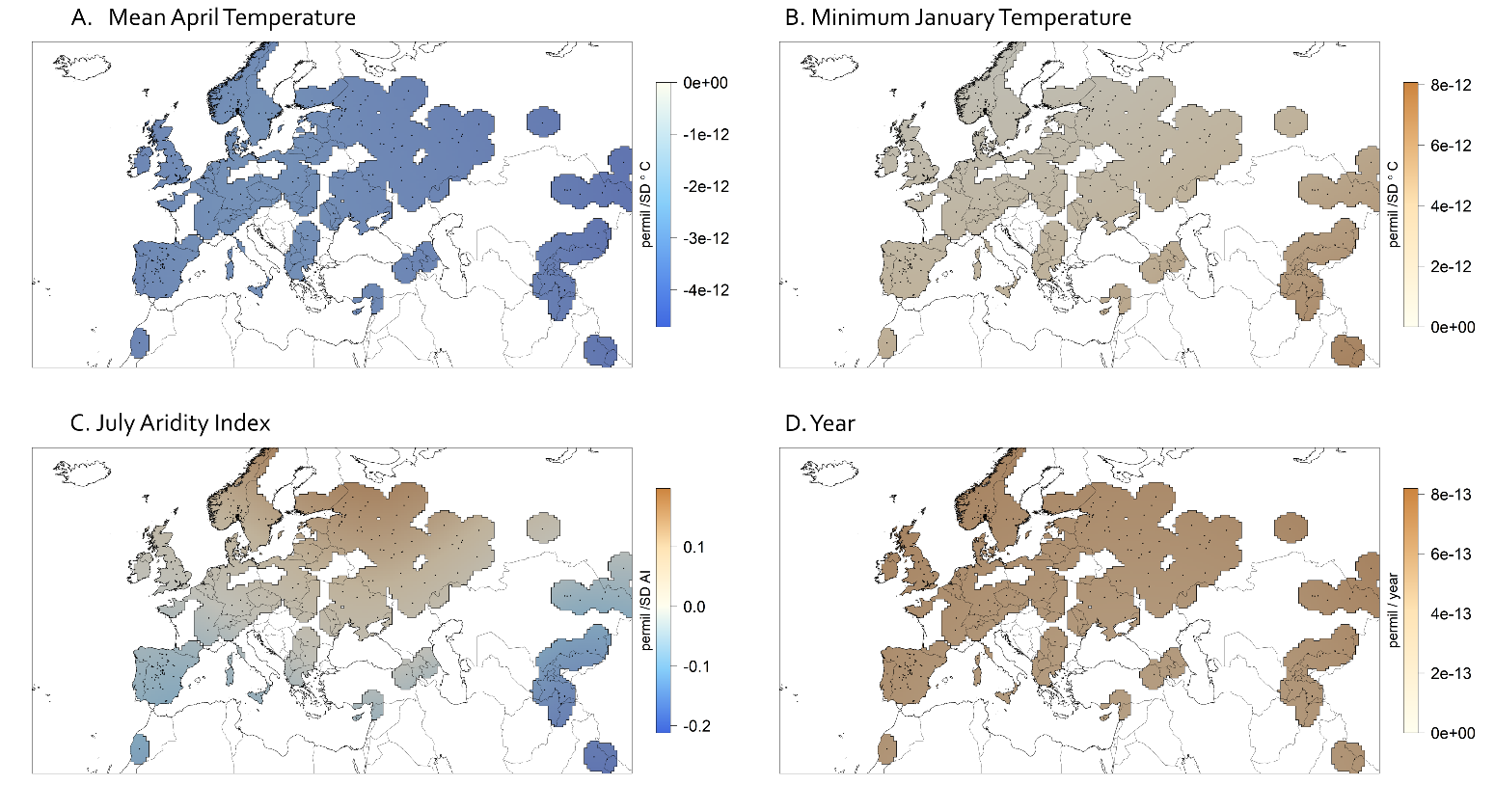
*Δ^13^C temporal anomaly model*

Figure S10: Coefficient surfaces of mean April temperature (A), minimum January temperature (B), July Aridity Index (C), and year (D) in a temporal anomaly model of Δ^13^C with a spatially varying intercept. For each climate variable, conditions in the year of collection are scaled to long term averages at that location. Shading represents areas where the 95% confidence interval of the estimated coefficent includes 0. Δ^13^C was not significantly related to any climate variable in this model.

Parametric coefficients:

Estimate	Std. Error	t value	Pr(>|t|)

(Intercept)	22.39182	0.06851	326.8	<2e-16

Approximate significance of smooth terms:

edf	Ref.df	F	p-value

s(lon,lat):intercept	3.586e+00	29	0.515	0.000633

s(lon,lat):tmn.1ano	4.001e-10	30	0.000	0.840415

s(lon,lat):tmp.4ano	4.838e-10	30	0.000	0.906028

s(lon,lat):ai.7ano	3.196e+00	30	0.133	0.266746

s(lon,lat):year2	4.415e-10	30	0.000	0.456981

R-sq.(adj) = 0.0453	Deviance explained = 6.17%

GCV = 1.8655 Scale est. = 1.829	n = 397

AIC = 1376.032

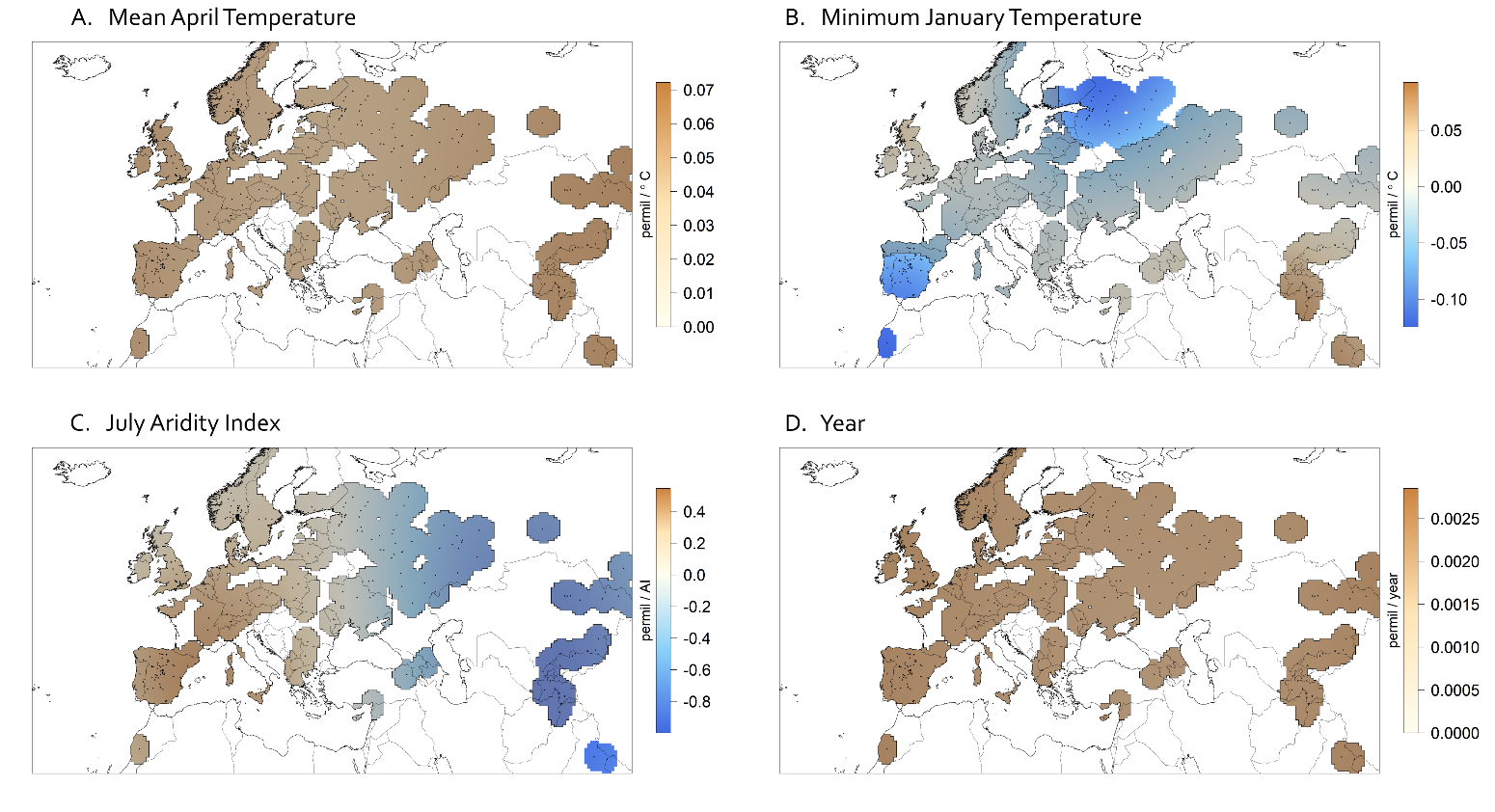
*Δ^13^C spatial model*

Figure S11: Coefficient surfaces of mean April temperature (A), minimum January temperature (B), July Aridity Index (C), and year (D) in a spatial model of Δ^13^C with a non-spatially varying intercept. For each climate variable, 50-year averages at each location are used. Shading represents areas where the 95% confidence interval of the estimated coefficent includes 0. Plants collected in areas of warmer winters of the Iberian Peninsula and western Russia have lower Δ^13^C values.

Parametric coefficients:

Estimate	Std. Error	t value	Pr(>|t|)

(Intercept)	22.3587	0.1086	205.9	<2e-16

Approximate significance of smooth terms:

edf	Ref.df	F	p-value

s(lon,lat):ltmn2	8.1369	29	1.047	8.05e-06

s(lon,lat):ltmp2	1.1489	29	0.131	0.03398

s(lon,lat):lai2	3.0431	29	0.315	0.00937

s(lon,lat):year2	0.8583	28	0.086	0.08016

R-sq.(adj) = 0.0768	Deviance explained = 10.4%

GCV = 1.7062 Scale est. = 1.6529	n = 454

AIC = 1532.514

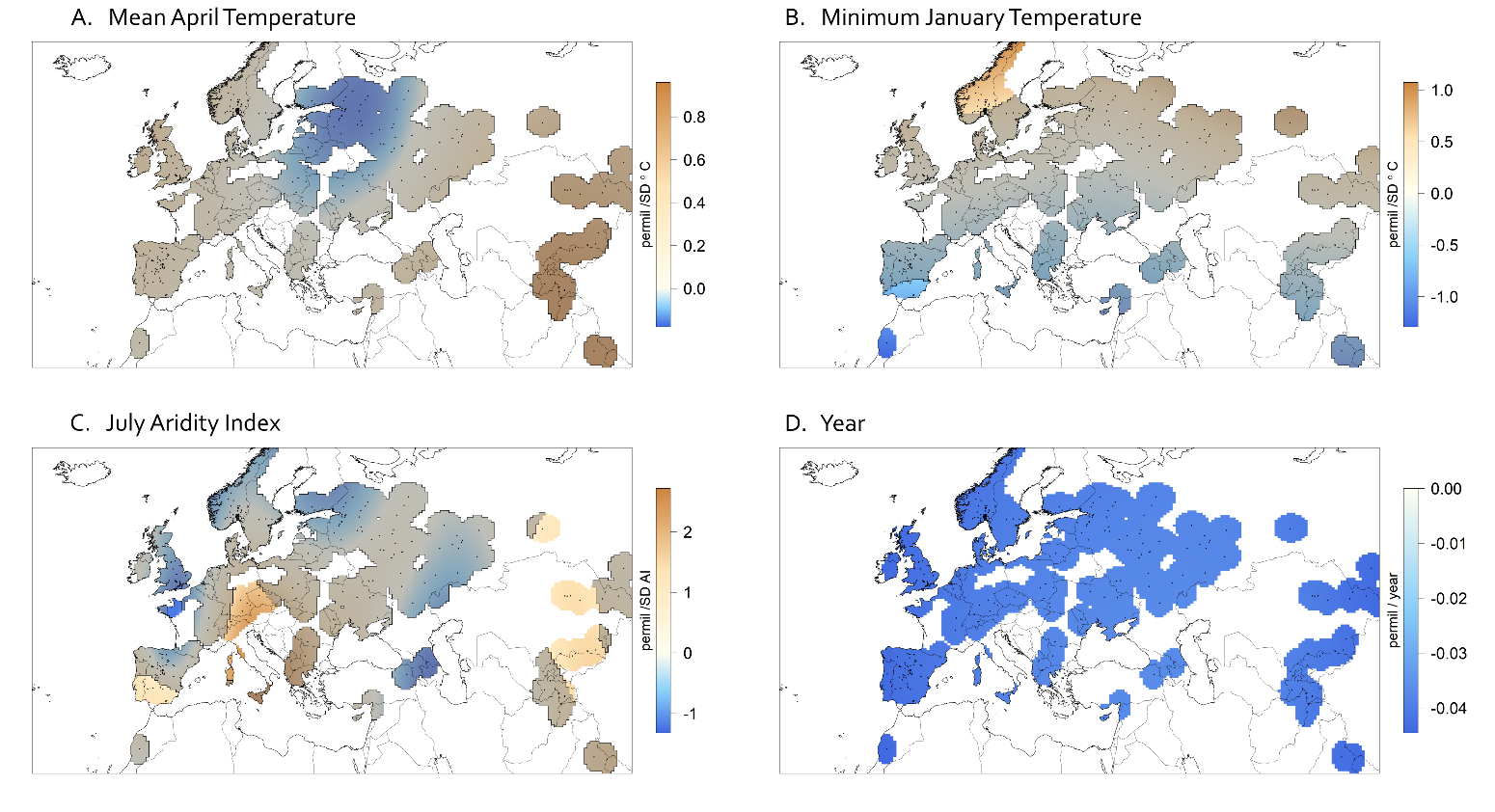
δ*^15^N temporal anomaly model*

Figure S12: Coefficient surfaces of mean April temperature (A), minimum January temperature (B), July Aridity Index (C), and year (D) in a temporal anomaly model of δ^15^N with a spatially varying intercept. For each climate variable, conditions in the year of collection are scaled to long term averages at that location. Shading represents areas where the 95% confidence interval of the estimated coefficent includes 0. δ^15^N increased in wetter years in a handful of regions. δ^15^N decreased through time across the entire range.

Parametric coefficients:

Estimate	Std. Error	t value	Pr(>|t|)

(Intercept)	4.1043	0.1613	25.44	<2e-16

Approximate significance of smooth terms:

edf	Ref.df	F	p-value

s(lon,lat):intercept	6.755	29	0.444	0.02792

s(lon,lat):tmn.1ano	4.028	30	0.399	0.00935

s(lon,lat):tmp.4ano	4.033	28	0.269	0.07161

s(lon,lat):ai.7ano	15.840	29	1.502	4.16e-05

s(lon,lat):year2	1.171	29	2.122	1.34e-15

R-sq.(adj) = 0.272	Deviance explained = 33.1%

GCV = 8.793 Scale est. = 8.0696	n = 399

AIC = 1998.863

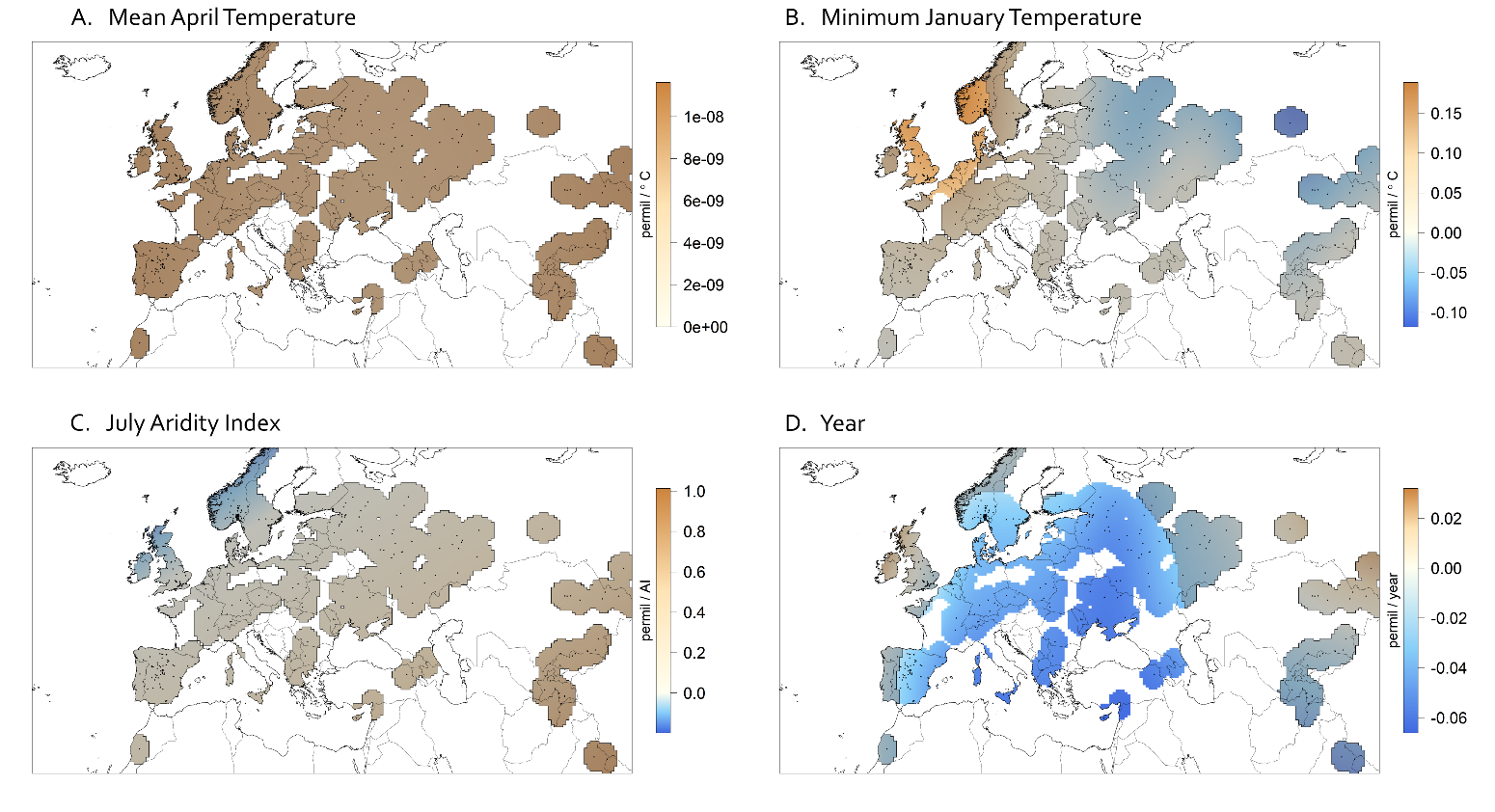
δ*^15^N spatial model*

Figure S13: Coefficient surfaces of mean April temperature (A), minimum January temperature (B), July Aridity Index (C), and year (D) in a spatial model of δ^15^N with a non-spatially varying intercept. For each climate variable, 50-year averages at each location are used. Shading represents areas where the 95% confidence interval of the estimated coefficent includes 0. As in the temporal model, δ^15^N decreases over time. Around the North Sea, plants collected in areas with warmer winters have higher δ^15^N values.

Parametric coefficients:

Estimate	Std. Error	t value	Pr(>|t|)

(Intercept)	3.8088	0.2055	18.54	<2e-16

Approximate significance of smooth terms:

edf	Ref.df	F	p-value

s(lon,lat):ltmn2	5.857e+00	29	0.463	0.0117

s(lon,lat):ltmp2	1.938e-07	30	0.000	0.3888

s(lon,lat):lai2	9.489e-01	30	0.085	0.0561

s(lon,lat):year2	9.760e+00	30	2.246	1.49e-12

R-sq.(adj) = 0.168	Deviance explained = 19.8%

GCV = 9.4682 Scale est. = 9.1058	n = 459

AIC = 2335.698

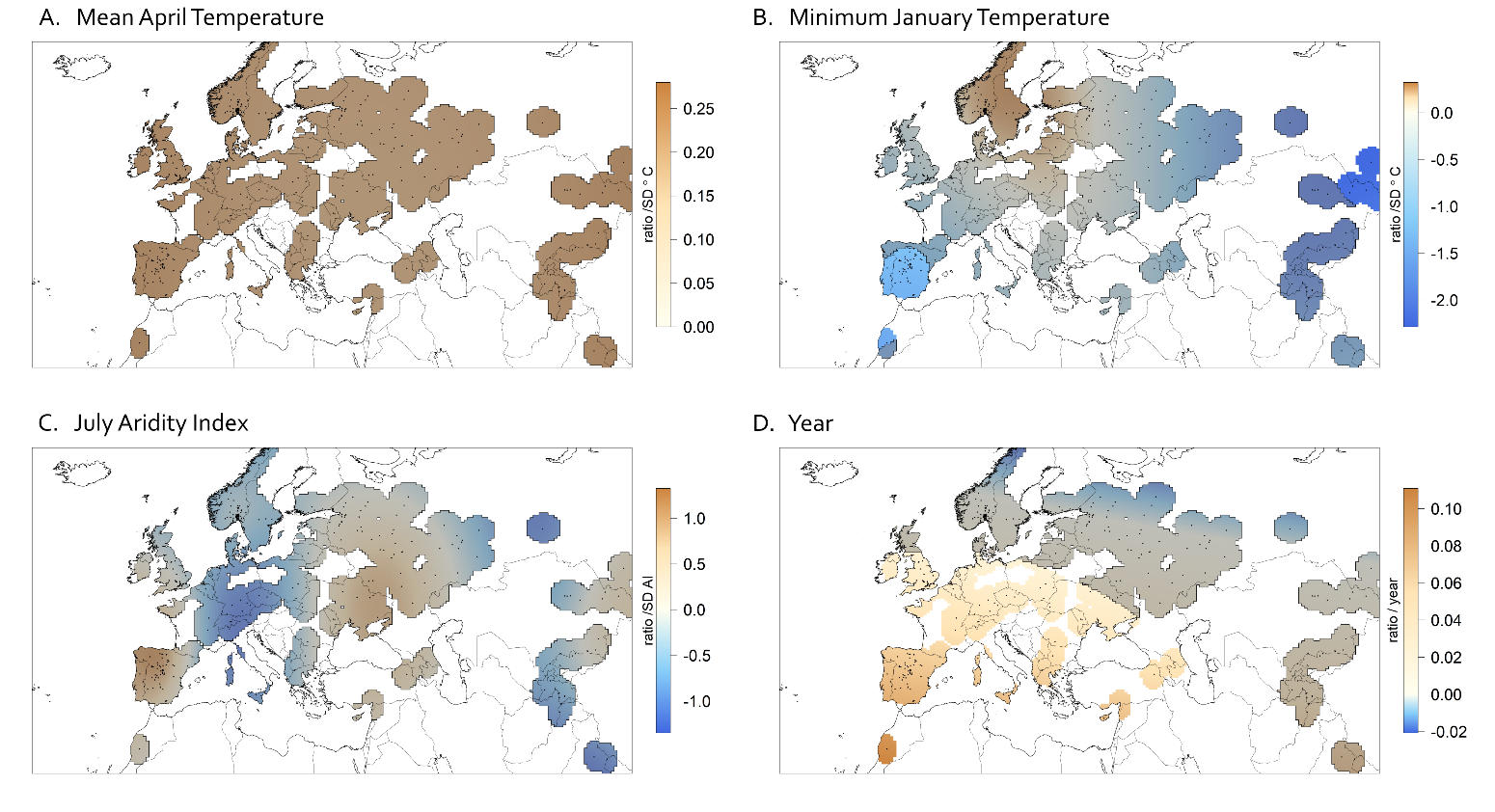
*C:N temporal anomaly model*

Figure S14: Coefficient surfaces of mean April temperature (A), minimum January temperature (B), July Aridity Index (C), and year (D) in a temporal anomaly model of C:N with a spatially varying intercept. For each climate variable, conditions in the year of collection are scaled to long term averages at that location. Shading represents areas where the 95% confidence interval of the estimated coefficent includes 0. C:N appears to be increasing in more recent years across much of southwestern Europe, while in the Iberian Peninsula and Northern Africa plants collected in years with warmer winters have lower C:N values.

Parametric coefficients:

Estimate	Std. Error	t value	Pr(>|t|)

(Intercept)	14.4549	0.3559	40.62	<2e-16

Approximate significance of smooth terms:

edf	Ref.df	F	p-value

s(lon,lat):intercept	10.1842	29	0.908	0.000564

s(lon,lat):tmn.1ano	3.7347	29	0.362	0.010930

s(lon,lat):tmp.4ano	0.4987	29	0.038	0.126287

s(lon,lat):ai.7ano	8.4939	29	0.319	0.307871

s(lon,lat):year2	2.2965	30	0.475 0	.000217

R-sq.(adj) = 0.186	Deviance explained = 23.7%

GCV = 38.75 Scale est. = 36.205	n = 399

AIC = 2591.706

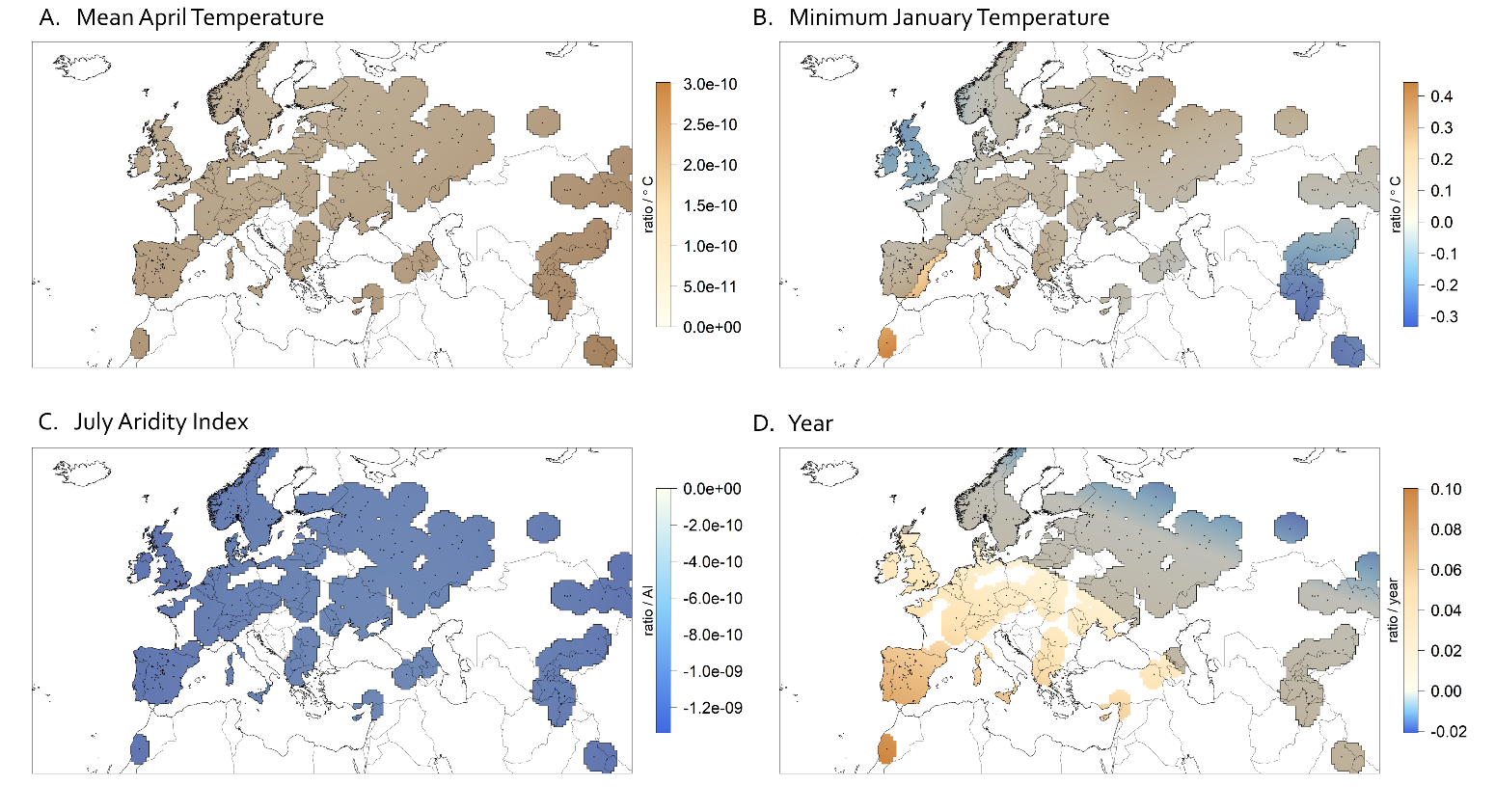
*C:N spatial model*

Figure S15: Coefficient surfaces of mean April temperature (A), minimum January temperature (B), July Aridity Index (C), and year (D) in a spatial model of C:N with a non-spatially varying intercept. For each climate variable, 50-year averages at each location are used. Shading represents areas where the 95% confidence interval of the estimated coefficent includes 0. As in the temporal model, C:N increases over time.

Parametric coefficients:

Estimate	Std. Error	t value	Pr(>|t|)

(Intercept)	14.5500	0.3889	37.41	<2e-16

Approximate significance of smooth terms:

edf	Ref.df	F	p-value

s(lon,lat):ltmn2	7.492e+00	29	0.780	0.000773

s(lon,lat):ltmp2	2.022e-09	30	0.000	0.326284

s(lon,lat):lai2	1.457e-09	30	0.000	0.273066

s(lon,lat):year2	2.433e+00	30	0.821	9.69e-07

R-sq.(adj) = 0.139	Deviance explained = 15.7%

GCV = 36.018 Scale est. = 35.159	n = 458

AIC = 2942.969

*Proportion N*. Note: the Hessian Matrix was not positive definite for these matrices, so the confidence intervals are unreliable.

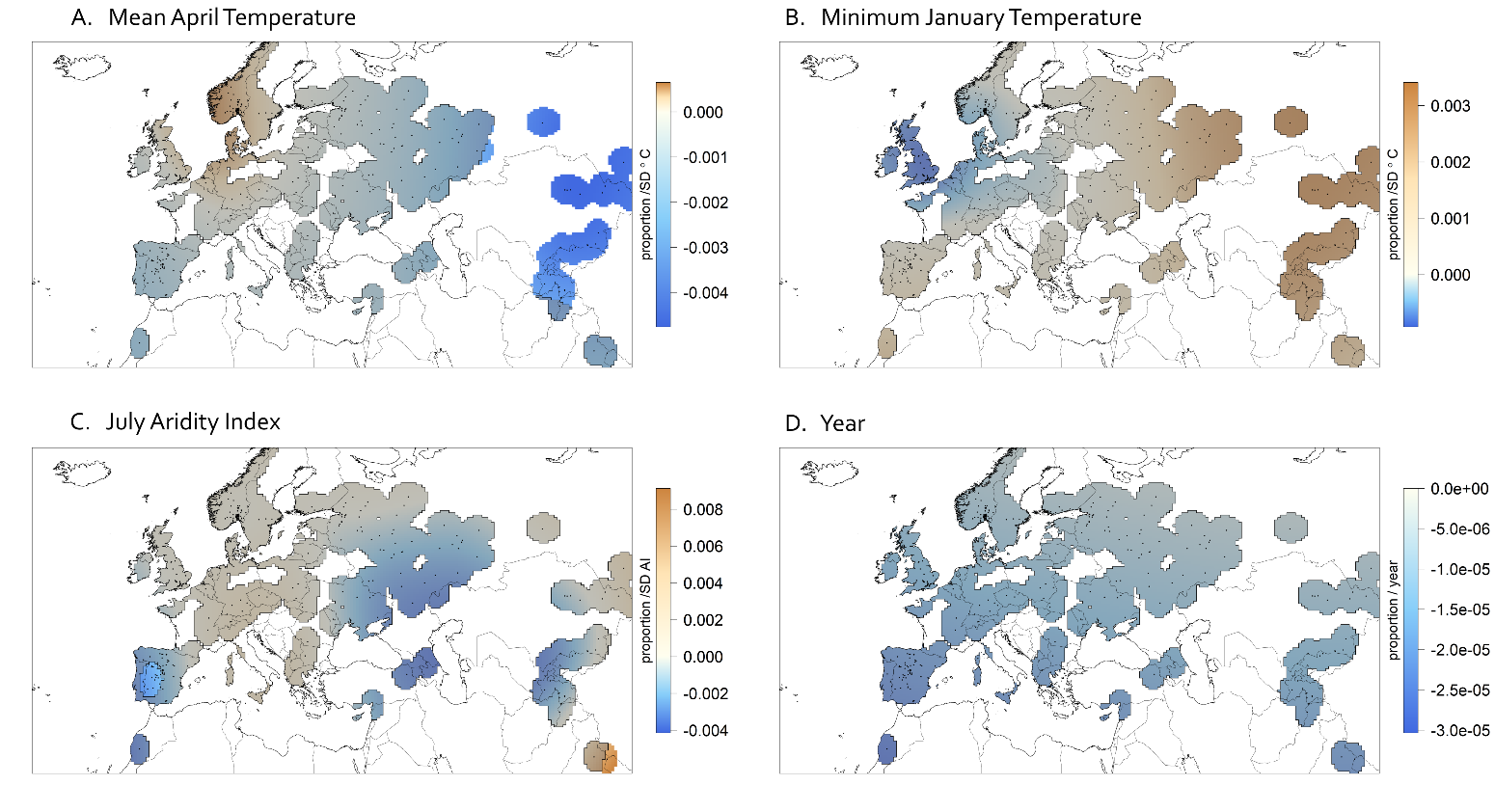
*Proportion N temporal anomaly model*

Figure S16: Coefficient surfaces of mean April temperature (A), minimum January temperature (B), July Aridity Index (C), and year (D) in a temporal anomaly model of Proportion N with a spatially varying intercept. For each climate variable, conditions in the year of collection are scaled to long term averages at that location. Shading is used to represent areas where the 95% confidence interval of the estimated coefficent includes 0. However, due to model limitations, the confidence intervals are unreliable.

Parametric coefficients:

Estimate	Std. Error	t value	Pr(>|t|)

(Intercept)	0.0279147	0.0004976	56.1	<2e-16

Approximate significance of smooth terms:

edf	Ref.df	F	p-value

s(lon,lat):intercept	10.5731	29	1.260	4.14e-06

s(lon,lat):tmn.1ano	5.0364	30	0.216	0.2078

s(lon,lat):tmp.4ano	4.0811	30	0.374	0.0100

s(lon,lat):ai.7ano	11.4126	30	0.708	0.0143

s(lon,lat):year2	0.7863	30	0.065	0.0557

R-sq.(adj) = 0.167	Deviance explained = 23.4%

GCV = 8.8158e-05 Scale est. = 8.0892e-05 n = 399

AIC = -2593.771

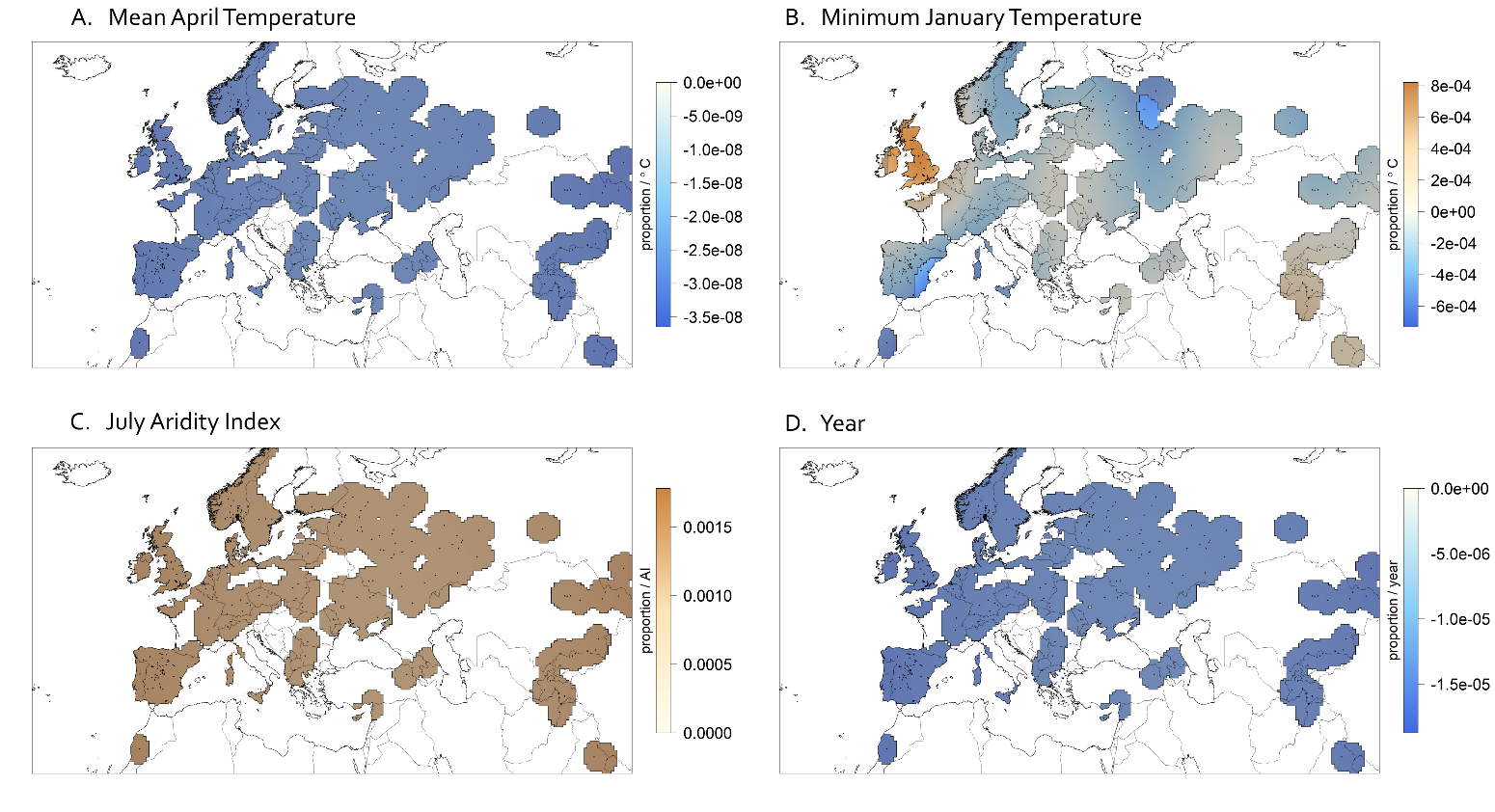
*Proportion N spatial model*

Figure S17: Coefficient surfaces of mean April temperature (A), minimum January temperature (B), July Aridity Index (C), and elevation (D) in a spatial model of Proportion N with a non-spatially varying intercept. For each climate variable, 50-year averages at each location are used. Shading is used to represent areas where the 95% confidence interval of the estimated coefficent includes 0. However, due to model limitations, the confidence intervals are unreliable.

Parametric coefficients:

Estimate	Std. Error	t value	Pr(>|t|)

(Intercept)	0.0271159	0.0006961	38.95	<2e-16

Approximate significance of smooth terms:

edf	Ref.df	F	p-value

s(lon,lat):ltmn2	1.213e+01	30	0.962	0.00164

s(lon,lat):ltmp2	2.179e-04	30	0.000	0.46093

s(lon,lat):lai2	6.863e-01	30	0.076	0.05431

s(lon,lat):year2	7.292e-01	30	0.076	0.06781

R-sq.(adj) = 0.0886	Deviance explained = 11.6%

GCV = 8.7838e-05 Scale est. = 8.5047e-05 n = 458

AIC = -2976.454

Environmental covariates removed from the final models:

The N deposition estimates we tested (Dentener, 2006) were spatially coarse and slightly correlated with year (r^2^ = 0.2). Elevation was not obviously correlated with year (r^2^ = 0.004), but the smooth term for elevation had an estimated concurvity greater than 0.9 in both the temporal and spatial models, which indicates that it could be approximated from the smooth terms of our other variables. When elevation and nitrogen deposition were included in the model as covariates, the Hessian matrices were not positive definite, and thus could not be used to obtain confidence intervals. To look at elevation, we tested temporal and spatial models that include elevation but not year.

Elevation tended to have a negative relationship with Δ^13^C as expected due to declining C_a_ at high elevation (Körner, Farquhar, & Roksandic, 1988). This was significant in the temporal climate anomaly model across much of Asia.

*Δ^13^C using elevation in place of year-temporal anomaly model*

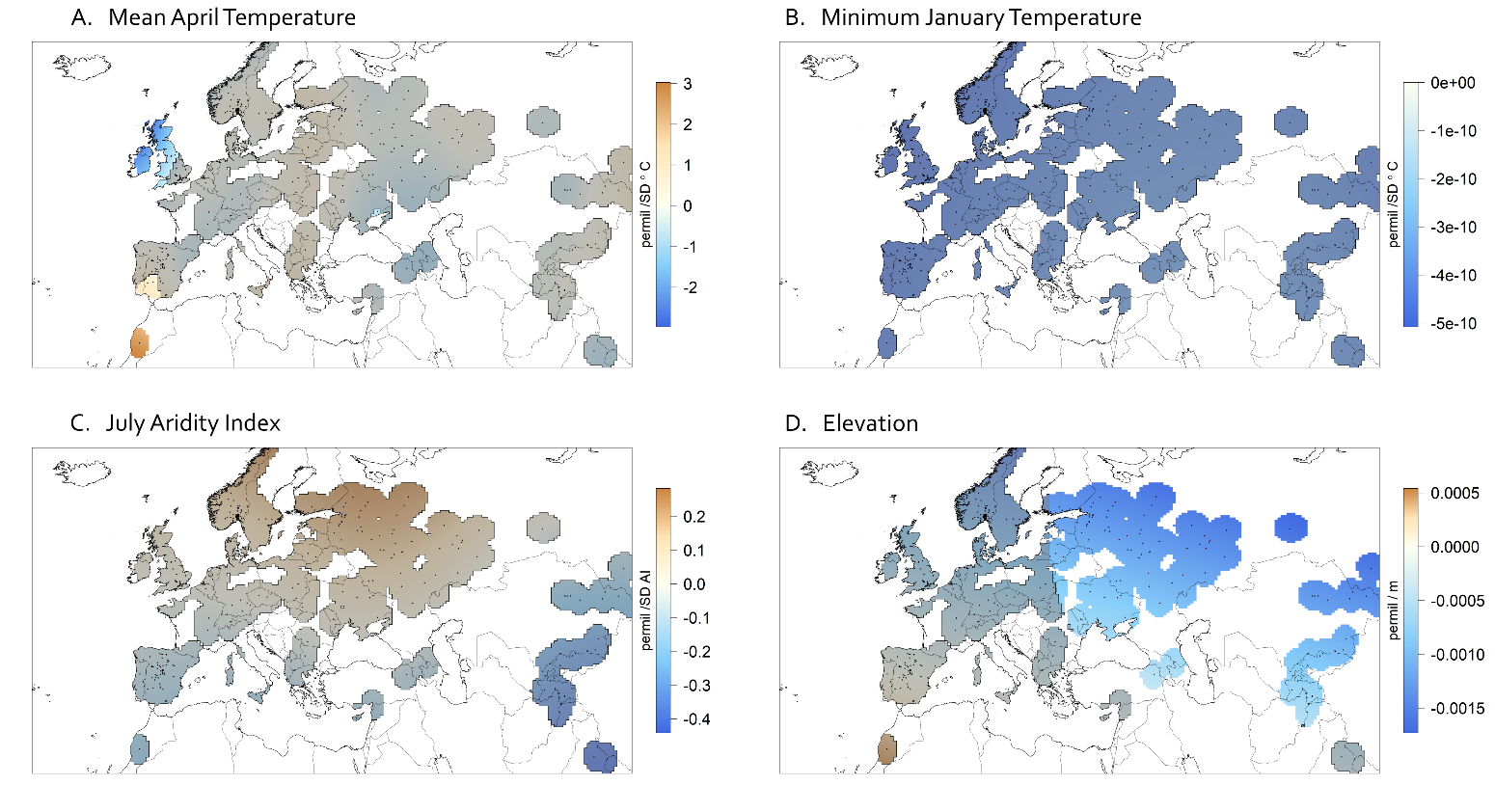
Figure S18: Coefficient surfaces of mean April temperature (A), minimum January temperature (B), July Aridity Index (C), and elevation (D) in a temporal anomaly model of Δ^13^C with a spatially varying intercept. For each climate variable, conditions in the year of collection are scaled to long term averages at that location. Shading represents areas where the 95% confidence interval of the estimated coefficent includes 0. Δ^13^C was significantly related to elevation, with plants collected from highter elevations having a lower Δ^13^C.

Parametric coefficients:

Estimate	Std. Error	t value	Pr(>|t|)

(Intercept)	22.32698	0.09459	236	<2e-16

Approximate significance of smooth terms:

edf	Ref.df	F	p-value

s(lon,lat):intercept	9.938e+00	29	0.837	0.00121

s(lon,lat):tmn.1ano	1.837e-08	30	0.000	0.72088

s(lon,lat):tmp.4ano	2.074e+01	30	1.115	0.02249

s(lon,lat):ai.7ano	3.984e+00	30	0.228	0.09089

s(lon,lat):ele2	2.904e+00	22	0.541	0.00274

R-sq.(adj) = 0.129	Deviance explained = 21.2%

GCV = 1.8483 Scale est. = 1.6687	n = 397

AIC = 1368.489

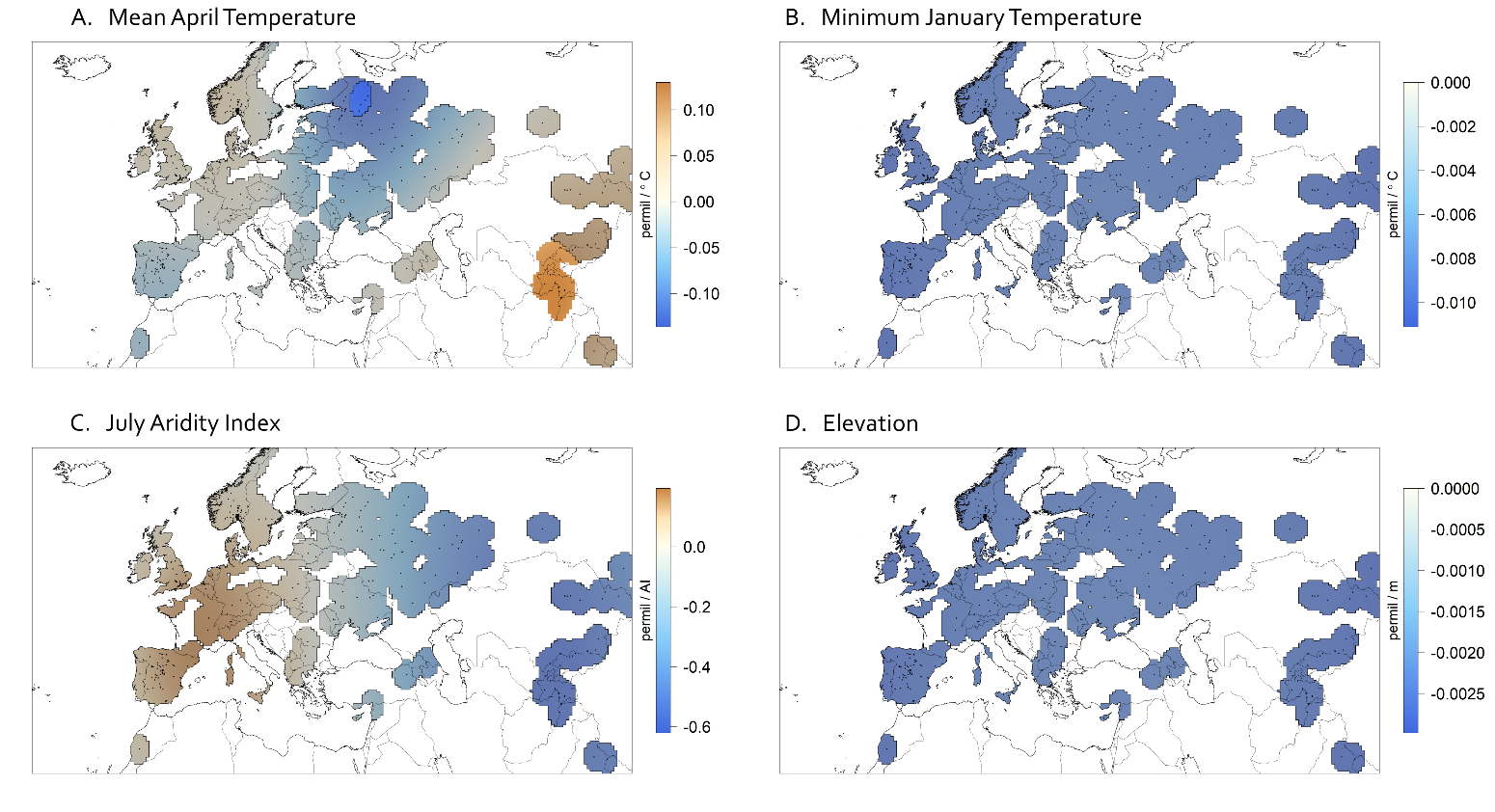
*Δ^13^C using elevation in place of year-spatial model*

Figure S19: Coefficient surfaces of mean April temperature (A), minimum January temperature (B), July Aridity Index (C), and elevation (D) in a spatial model of Δ^13^C with a non-spatially varying intercept. For each climate variable, 50-year averages at each location are used. Shading represents areas where the 95% confidence interval of the estimated coefficent includes 0. Plants collected in areas of warmer springs in Central Asia may have higher Δ^13^C values while plants collected in areas of warmer springs northwestern Asia may have lower Δ^13^C values.

Parametric coefficients:

Estimate	Std. Error	t value	Pr(>|t|)

(Intercept)	22.37853	0.08889	251.8	<2e-16

Approximate significance of smooth terms:

edf	Ref.df	F	p-value

s(lon,lat):ltmn2	0.4356	29	0.034	0.103469

s(lon,lat):ltmp2	5.4701	29	0.676	0.000192

s(lon,lat):lai2	2.1325	29	0.135	0.092943

s(lon,lat):ele2	0.7811	29	0.138	0.013542

R-sq.(adj) = 0.0675	Deviance explained = 8.57%

GCV = 1.7064 Scale est. = 1.6695	n = 454

AIC = 1532.786

*Δ^13^C using only samples collected before 1950.* Rising atmospheric CO_2_ concentrations may mask climate associations (Drake et al., 2017). However, Δ^13^C before 1950 is still not significantly related to climate for the temporal model or the year only model, and the spatial model indicates a negative association between Δ^13^C and aridity in the eastern part of the native range.

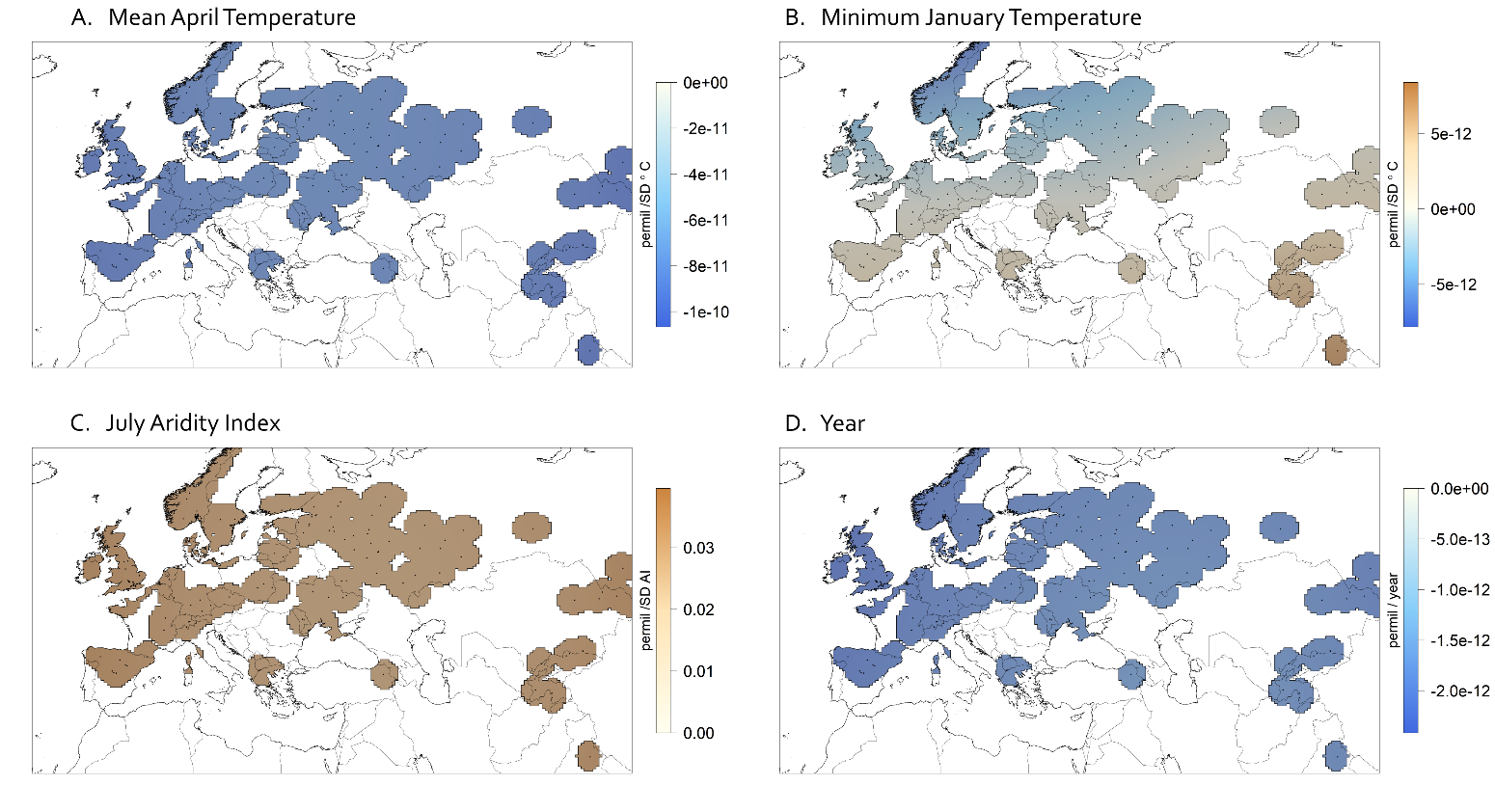
*<1950s temporal anomaly model*

Figure S20: Coefficient surfaces of mean April temperature (A), minimum January temperature (B), July Aridity Index (C), and elevation (D) in a temporal model of Δ^13^C with a spatially varying intercept. Only samples before 1950 were included. For each climate variable, conditions in the year of collection are scaled to long term averages at that location. Shading represents areas where the 95% confidence interval of the estimated coefficent includes 0. Δ^13^C was not significantly related to any covariate.

Parametric coefficients:

Estimate	Std. Error	t value	Pr(>|t|)

(Intercept)	22.43826	0.08836	254	<2e-16

Approximate significance of smooth terms:

edf	Ref.df	F	p-value

s(lon,lat):intercept	1.557e+01	29	1.282	0.00113

s(lon,lat):tmn.1ano	2.495e-09	30	0.000	1.00000

s(lon,lat):tmp.4ano	2.062e-09	30	0.000	0.65451

s(lon,lat):ai.7ano	3.128e-01	28	0.015	0.23257

s(lon,lat):year2	3.517e-10	30	0.000	0.37998

R-sq.(adj) = 0.154	Deviance explained = 23.1%

GCV = 1.5191 Scale est. = 1.3734	n = 176

AIC = 573.3227

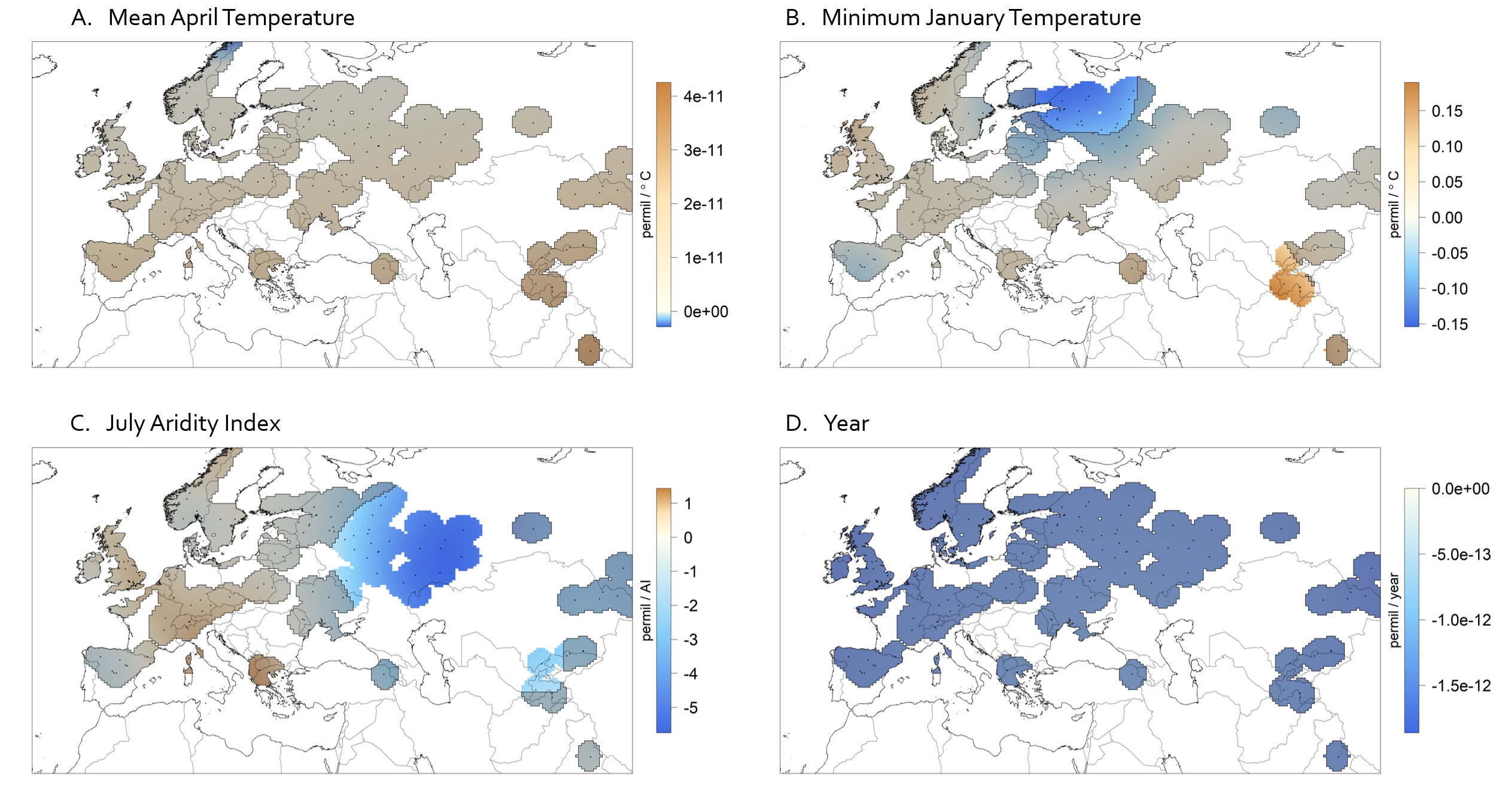
*<1950s spatial model*

Figure S21: Coefficient surfaces of mean April temperature (A), minimum January temperature (B), July Aridity Index (C), and elevation (D) in a spatial model of Δ^13^C with a non-spatially varying intercept. Only samples collected before 1950 were included. For each climate variable, 50-year averages at each location are used. Shading represents areas where the 95% confidence interval of the estimated coefficent includes 0. Plants collected in years of warmer winters in Central Asia may have higher Δ^13^C values while plants collected in warmer winters northeastern Europe may have lower Δ^13^C values.

Parametric coefficients:

Estimate	Std. Error	t value	Pr(>|t|)

(Intercept)	22.1467	0.1491	148.6	<2e-16

Approximate significance of smooth terms:

edf	Ref.df	F	p-value

s(lon,lat):ltmn2	1.077e+01	29	1.059	8.12e-05

s(lon,lat):ltmp2	5.504e-10	30	0.000	0.184652

s(lon,lat):lai2	8.046e+00	30	0.749	0.000653

s(lon,lat):year2	1.455e-09	30	0.000	0.798388

R-sq.(adj) = 0.147	Deviance explained = 21.6%

GCV = 1.3584	Scale est. = 1.2424	n = 232

AIC = 729.6557
